# Loop extrusion creates rare, long-lived encounters underlying enhancer-promoter communication

**DOI:** 10.1101/2025.09.24.678119

**Authors:** Mattia Ubertini, Nessim Louafi, Kristina Landry, Pavel I. Kos, Gregory Roth, Edoardo Marchi, Julie Cramard, Guido Tiana, Luca Giorgetti

**Author notes:** equal contribution (Mattia Ubertini, Nessim Louafi).

## Abstract

Enhancers regulate transcription from distal genomic positions, but how their spatial encounters with promoters drive activation remains unclear. Using polymer simulations and high-resolution live-cell microscopy, we identify rare but long-lived chromatin encounters arising from cohesin-mediated loop extrusion. These events occur when cohesin loads near the midpoint between two loci and extrudes them through a defined spatial radius, producing encounter durations that exceed those of random collisions. We show that such encounters explain observed nonlinear relationships between contact probability and transcription, and accurately predict transcriptional changes upon perturbation of cohesin or its cofactors. Our findings support a time-gated model of distal enhancer-promoter communication in which only rare, long-lived and mostly extrusion-driven encounters are productive, offering a unifying framework for how chromosome dynamics control transcription in single cells.

## Introduction

The three-dimensional (3D) folding of chromosomes plays a central role in transcriptional regulation in mammalian cells, notably by modulating the probabilities that enhancers and promoters come into spatial proximity (*1*). Such encounters are thought to enable the transmission of regulatory information through molecular interactions involving transcription factors, cofactors, and components of the transcriptional machinery, ultimately licensing transcriptionally competent RNA polymerase II (Pol II) at the promoter. However, the mechanistic details governing these interactions remain incomplete, including the sequence of molecular events, the identity of participating factors, and the spatial constraints under which regulation occurs. It remains unclear whether enhancer-promoter communication depends on direct physical contact between proteins bound to DNA and their cofactors, or on longer-range interactions mediated by higher-order biomolecular assemblies (*2*). Uncertainty also persists over whether transcriptional activation requires only brief encounters or stable proximity, and how this relates to transcription factor binding, assembly of regulatory complexes, and the kinetics of biochemical reactions at regulatory loci (*3*). These open questions limit our understanding of how enhancer-promoter encounters are converted into transcriptional output at the single-cell level.

In mammals, chromosome folding is orchestrated by the ATP-dependent loop extrusion activity of the cohesin complex, which is itself controlled by regulators of cohesin loading rate, extrusion velocity (such as NIPBL) (*4*) or lifetime on DNA (such as WAPL) (*5*), and is halted when cohesin encounters CTCF bound to DNA in a defined orientation (*6*). The resulting formation of CTCF-anchored loops partitions the genome into domains of preferential physical interactions (*7*, *8*) that are essential for proper enhancer-promoter communication and whose disruption can be causal to genetic disease (*9*). Recent studies have also clarified that cohesin’s loop extrusion activity *per se*, even without stalling by CTCF sites, is crucial for transcriptional regulation. Acute depletion of the cohesin subunit RAD21 or its loading and elongation factor NIPBL revealed that loop extrusion is necessary for enhancers to control promoters when their genomic separation exceeds few tens of kilobases (kb) (*10–13*). Cohesin appears on the other hand to be dispensable either when an enhancer is located closer to a promoter (*10–12*), or in the presence of additional promoter-proximal weak enhancers, which can buffer the effects of cohesin depletion (*13*). Yet, the mechanistic bases of how encounters mediated by loop extrusion are translated into transcriptional events are not understood. It is also unclear to what extent loop extrusion is responsible for the recently observed nonlinear relationships of promoter transcription levels as a function of genomic separation and contact probability with an enhancer (*10*, *14–17*), and whether or not it underlies the observation that mammalian enhancers mostly control the frequency of transcriptional bursts (*18–20*).

While the structural effects of loop extrusion and its regulation by cohesin cofactors and CTCF are increasingly well characterized, far less is known about how these factors shape the dynamics of chromatin folding. In particular, it remains unclear how loop extrusion influences the frequency and duration of spatial encounters between enhancers and promoters, and whether such dynamics govern transcriptional activity at promoters in single cells. Recent live-cell imaging studies have provided the first quantitative measurements of the lifetimes of loops anchored by convergent CTCF sites, revealing durations of approximately 10 to 30 minutes and frequencies that scale inversely with genomic distance between the anchor sites (*21–23*). However, substantial uncertainty persists regarding the frequency and duration of physical encounters mediated by cohesin when not constrained by CTCF. Our previous work, as well as recent ultra-high-resolution live imaging studies, suggests that in the absence of CTCF encounters between genomic loci are short-lived (*21*, *24*), and that loop extrusion accelerates the recurrence of encounters (*21*, *24*, *25*). Yet, quantitative estimates of the duration and frequency of such transient events in dual-color imaging of pairs of genomic loci are highly sensitive to the choice of spatial thresholds for defining proximity. Even more critically, they are severely affected by the localization error inherent to sub-diffraction fluorescence microscopy which inevitably leads to the detection of artifactual encounter events (*26*). As a result, reported estimates of encounter durations range widely - from several minutes within a 150-nm range (*21*) to one second or less within a 50-nm threshold (*24*).

Uncertainty about the duration and frequency of physical encounters along chromosomes also reflects our limited understanding of the timescales of looping events in polymer systems. Theoretical studies in polymer dynamics have primarily focused on estimating the waiting times between consecutive encounters of monomer pairs in simplified polymer models, a problem that is closely related to cyclization and reaction kinetics in polymer chemistry (*27*, *28*). In contrast, there are virtually no theoretical predictions for how long two monomers remain within a defined encounter radius outside basic polymer models (*29*), let alone in systems subject to active processes such as loop extrusion. Thus, there is currently no theoretical framework that is able to provide realistic expectations for the frequency and duration of chromosomal encounters in the presence of loop extrusion, how they are distributed in time and across cells, and how they could be related to the transmission of regulatory information between enhancers and promoters.

Here, we address these unresolved questions through a combination of polymer simulations, live-cell imaging of chromosome dynamics at exceptionally high spatiotemporal resolution, and genome engineering of mouse embryonic stem cells (mESC). Using polymer models that recapitulate both the statistical and dynamic properties of chromatin, we find that in the absence of loop extrusion, all encounters (even those occurring within hundreds of nanometers) are extremely transient, rarely lasting more than a few seconds. In contrast, we identify a distinct class of rare, long-lived encounters that arise specifically from loop extrusion, and even in the absence of CTCF roadblocks. These events occur only when cohesin loads near the midpoint between two loci, extruding them first into, and then out of, the encounter radius, resulting in contact durations of tens of seconds, which far exceed those of random polymer collisions and scale inversely with the extrusion velocity. While loop extrusion also increases the frequency of encounters, we show that this effect results from enhanced compaction and does not constitute a distinct dynamic signature. Instead, the emergence of rare, prolonged contacts represents the only specific temporal hallmark of active loop extrusion.

We further show that, although finite sampling rates and localization error in live-cell microscopy preclude precise determination of absolute encounter durations, the dynamic signature of loop extrusion can still be observed by measuring ‘apparent’ encounter times under strictly calibrated experimental conditions with exceptionally high spatiotemporal resolution. We demonstrate that this resolution can be achieved through improvements to widefield live-cell fluorescence microscopy, enabling imaging of two fluorescently labeled chromosomal loci in mESCs at 2-second intervals with spatial uncertainty reduced to a few tens of nanometers - which is comparable to the physical scale at which regulatory interactions might occur at enhancer–promoter interfaces (*30*). By comparing the dynamics of these loci before and after inducible depletion of RAD21 and to the dynamics of loci separated by a different genomic distance, we validate model predictions and provide experimental evidence for the existence of rare, long-lived encounters mediated by cohesin-driven loop extrusion.

Finally, we show that the simple hypothesis that an enhancer exerts its regulatory effect only through encounters exceeding a minimum duration (thereby favoring rare, loop-extrusion mediated events) accurately predicts previous quantitative measurements of transcriptional output as a function of enhancer-promoter distance in the absence of CTCF (*15*), as well as the transcriptional effects of RAD21 and NIPBL depletion on distance-dependence transcription levels (*10–13*). This model further implies that although an enhancer may encounter its target promoter many times during the cell cycle, most of these encounters are unproductive and only a small subset of events should lead to actual transmission of regulatory information to the promoter. We predict such productive encounters to occur very rarely, from less than once to a few times per cell cycle, depending on the encounter radius required for promoter regulation and enhancer-promoter genomic separation.

By revealing the existence of rare but long-lived encounters mediated by cohesin, and linking such encounters to transcriptional activation through the minimal assumption that enhancers activate promoters *via* time-gated molecular processes, our work provides a unified explicative framework for understanding how chromosome structure and dynamics govern transcriptional regulation in mammalian cells.

## Results

### Loop extrusion dynamics generates rare but long-lived encounters between pairs of loci

To establish a framework for investigating the frequency and duration of physical encounters between genomic loci and the specific contributions of loop extrusion, we first performed simulations using a coarse-grained polymer model that captures the static and dynamic properties of the chromatin fiber that were experimentally observed in mammalian cells both in the presence and absence of cohesin. Widely used open-chain polymer models of the chromatin fiber (*21*, *22*) fail to reproduce the observed inverse power-law exponent (∼ −1) of contact probability as a function of genomic distance in Hi-C and Micro-C experiments after depletion of cohesin (**Fig. 1A**) (*31*). We thus turned to ring polymer models with excluded volume, whose topological constraints make them capable of reproducing such scaling (*32*). Specifically, we represented four megabases of chromatin as a ring polymer at a volume fraction of 30%, with each monomer corresponding to 1 kilobase of DNA, and analyzed segments of up to 1 Mb each to avoid artefactual topological effects due to the closure of the polymer (see Methods). As expected from polymer theory (*33*, *34*), such model also recapitulates the subdiffusive behavior of the mean-square displacement of the vector distance between two genomic loci, as measured in this work (**Supplementary Fig. 1A**; see results below) and in recent studies (*21*, *24*, *25*). This ring polymer is thus a useful effective model for the statistical and dynamic properties of the chromatin fiber without loop extrusion. We then incorporated loop extrusion in this model by simulating bidirectional extruders (Methods). Unless differently stated, we employed parameters that best matched experimentally measured contact probabilities observed in wild-type mESC (*31*) (**Fig. 1B**) (average extruder lifetime: 10 minutes, average velocity: 0.3 kb/second (s), mean separation between extruders: 150 kb resulting in an average loop size of approximately 115 kb), which were in the range of available experimental estimates (*35*, *36*). Time and spatial units were estimated by matching the mean squared displacement and average physical distances measured in live mESC for two loci separated by 150kb (see Methods and **Fig. 2** below for the experimental setup).

**Figure 1:**
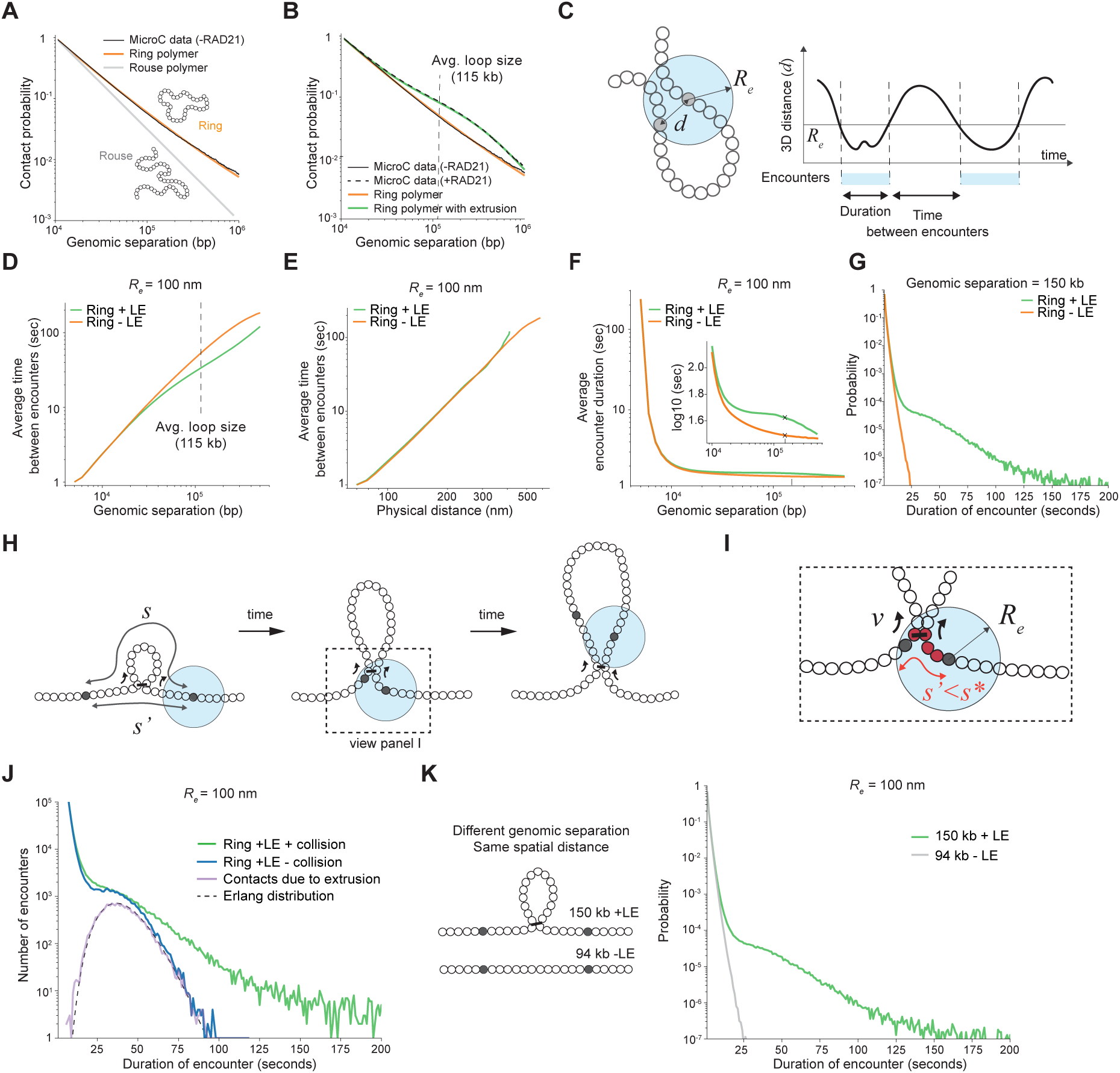
Polymer physics predicts the existence of long-lived encounters mediated by loop extrusion. **A.** Contact probability as a function of genomic separation in a Rouse polymer, the ring polymer used in this study and experimental MicroC data in the absence of RAD21 (experimental data from Ref. (*31*)). **B.** Contact probability as a function of genomic separation for the ring polymer without or with loop extrusion (LE) compared to MicroC data in the absence or presence of RAD21 (data from Ref. (*31*)). **C.** Schematics of definitions of encounter duration and time between consecutive encounters. Encounters are defined as instances where the 3D physical distance between two monomers (*d)* is smaller than an arbitrary encounter radius 𝑅_𝑒_. **D.** Average time between encounters as a function of genomic separation between monomers in a ring polymer with or without loop extrusion. Dotted line: effective average loop size computed from polymer simulations. **E.** Same as D but as a function of the average physical distance between monomers. **F.** Average duration of encounter in a sphere of radius 100 nm as a function of the genomic separation between monomers with and without loop extrusion. Inset: magnification for distances >10kb. Crosses indicate the genomic separation of 150 kb, see panel G. **G.** Distribution of encounter durations between monomers separated by 150 kb. **H.** Illustration of the origin of encounters mediated by loop extrusion. 𝑠: genomic separation between monomers, 𝑠′: effective separation between monomers in the presence of a loop, 𝑣: extrusion speed, 𝑠^∗^: separation for which the pair of monomers is always inside the sphere of radius 𝑅_𝑒_. **J.** Number of encounters as a function of their duration for models allowing or preventing extruders to bypass each other. Purple line: Number of encounters due to loop-extrusion (defined in simulations as those where an extruder links the two monomers or any of the two flanking monomers). Dotted line: analytical prediction reporting the Erlang distribution for the corresponding extrusion parameters. **K.** Distributions of encounter durations for two pairs of monomers separated by 150 kb and 94 kb in the polymer model, displaying equal average physical distance with and without extrusion, respectively.

**Figure 2:**
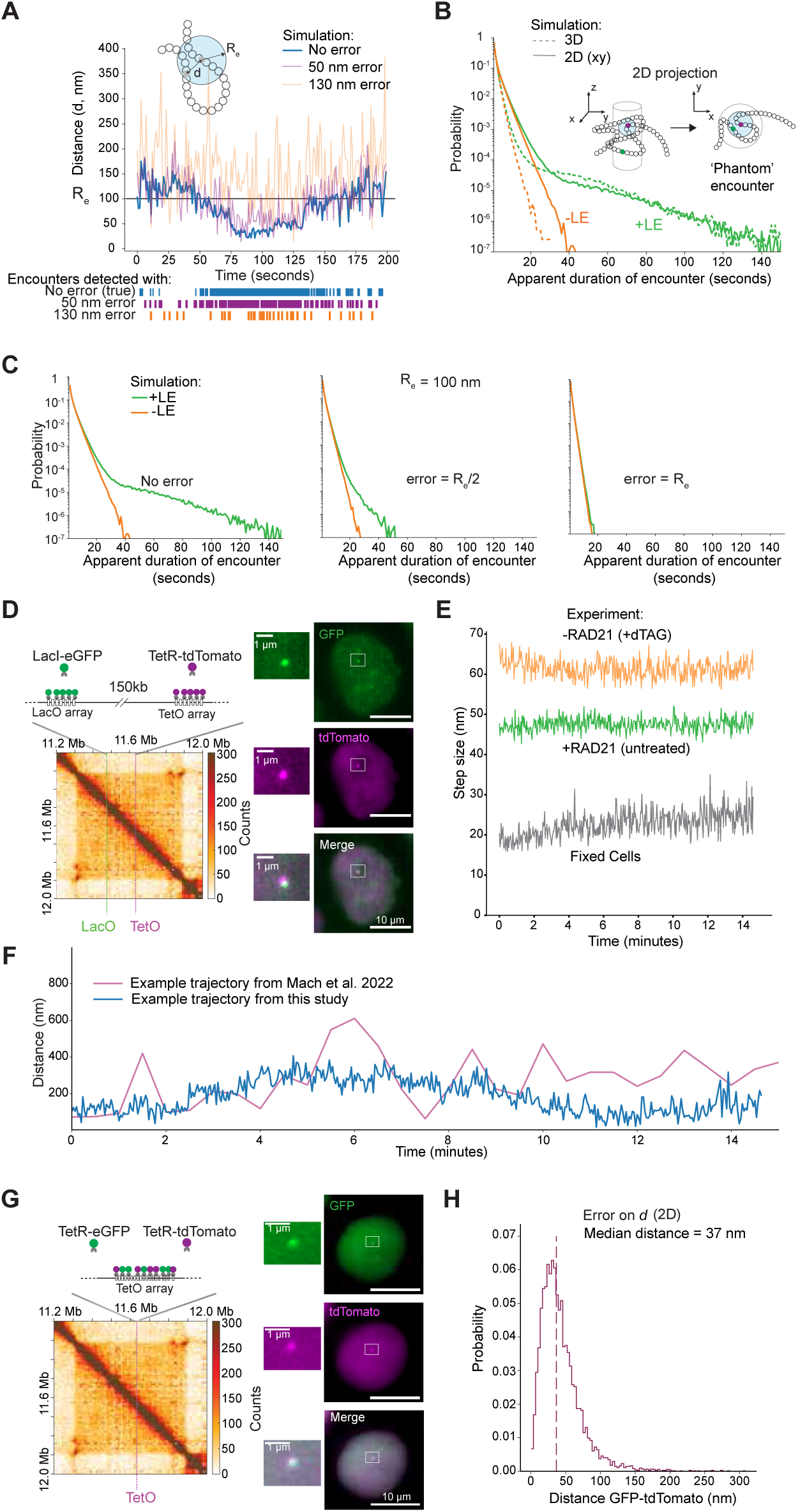
Experimental conditions enabling detection of encounters mediated by loop extrusion. **A.** Illustration of the effect of experimental error on measurement of encounter durations. A representative trajectory from polymer simulations is compared with versions of the same trajectory with overlaid 3D Gaussian error of 50 and 130 nm on average. **B.** Distributions of encounter durations before and after projection onto the xy plane in simulations with and without loop extrusion. The inset scheme illustrates z-projection effects and phantom encounters. **C.** Distributions of apparent encounter durations for increasing levels of simulated 2D error on distances. From left to right: the error is 0, the error is half of the encounter radius, the error equals the encounter radius. **D.** Left: Schematics of experimental systems. Top: insertion of TetO and LacO arrays separated by 150 kb within a ‘neutral’ TAD on chromosome 15 in mESCs. Arrays are visualized by binding of LacI**-eGFP and TetR-tdTomato, respectively. Bottom, tiled Capture-C map (6.4-kb resolution) in mESCs in a region of 2.6 Mb surrounding the engineered TAD. Dashed lines, genomic positions of LacO and TetO insertions. Capture-C map from Ref. (*21*). Right: representative fluorescence microscopy images of mESCs in the green (LacI**-eGFP) and red (TetR-tdTomato) channels, and the two channels overlaid (exposure time 100 ms, maximum intensity projection). **E.** Average frame-to-frame 2D (xy) displacement over time in live-cell movies, for cells treated with dTAG-13 for RAD21 depletion (orange curve, 780 tracks) and untreated cells (green curve, 626 tracks). Fixed cells are shown as a gray curve (120 tracks; see Methods)**. F.** Representative time-resolved LacO-TetO distances following optimization of microscope setup in this study and from a previous study using the same cells (*21*). **G.** Left: Schematics of control mESC line with a single TetO array bound by both TetR-eGFP and TetR-tdTomato. Right: as in panel **D** but for the control mESC line. **H.** Distributions of 2D distances between TetR-eGFP and TetR-tdTomato signals in the control cell line described in panel G. TetR-eGFP and TetR-tdTomato signals were filtered to match the signal-to-background ratios observed in the cell line described in panel D (see Methods). Dotted line: median value.

We next operationally defined encounters as events where two monomers simultaneously reside within a sphere of radius 𝑅_𝑒_centered on either monomer. Because the spatial range required for effective enhancer–promoter communication remains unknown, we systematically varied 𝑅_𝑒_ across a broad range (from 25 to 200 nm, corresponding to approximately the distance between 2 to 40 monomers), and for each radius characterized both the waiting times between consecutive encounters and the durations of individual encounters (**Fig. 1C**). In the absence of loop extrusion, the average waiting time between consecutive encounters increased with genomic separation, broadly following a power-law relationship as recently observed (*24*, *25*). Incorporating loop extrusion markedly reduced waiting times at genomic separations close to the average extruded loop size (∼115 kb), indicating that extrusion increases the frequency of encounters between monomers (**Fig. 1D**), independently of the chosen encounter radius (**Supplementary Fig. 1B**). These predictions align with recent experimental observations reporting increased encounter frequencies across multiple spatial scales in the presence of cohesin (*21*, *24*, *25*). However, the reduction in waiting times induced by extrusion is exclusively due to increased polymer compaction: when we plotted waiting times against average 3D spatial distance between monomers rather than their genomic separation, the curves with and without loop extrusion collapsed onto the same power-law behavior (**Fig. 1E**). Thus, while loop extrusion accelerates search kinetics by compacting the polymer, this acceleration does not represent a direct dynamic signature unique to the extrusion process itself. Finally, for genomic separations of ∼100–150 kb, which are typical of E-P separations, the predicted encounter times fall within the tens of seconds range, consistent with recent estimates (24), and occur significantly more frequently than the inter-burst intervals recently reported by (18).

We next examined the durations of encounters, for which no prior theoretical expectations exist beyond the ideal chain (*29*). We found that at short genomic separations, such that monomers reside most of the time inside 𝑅_𝑒_, encounter durations are longer than at longer genomic separations, where physical distances become significantly larger than 𝑅_𝑒_ (**Fig 1F**). Loop extrusion produced only a modest increase in average encounter duration in the genomic range of the average extruded loop (∼115 kb) (**Fig. 1F**, inset). This is due to the fact that the vast majority of physical encounters arise from thermal fluctuations of the fiber and are very transient, both with and without loop extrusion (**Fig. 1G**). Crucially, however, loop extrusion gives rise to a distinct class of rare, long-lived contacts persisting for tens of seconds, which were entirely absent without extruders (**Fig. 1G**). We note that these extended interactions do not result from extruders that are stalled at architectural barriers such as CTCF sites, as there were none in the simulations. Introducing impermeable extrusion barriers led indeed to the formation of contacts whose lifetime is the same as the residence time of the extruders themselves (10 min in these simulations) (**Supplementary Fig. 1C**).

The complete absence of long-lived encounters in simulations lacking extruders suggests that these events arise specifically from the loop extrusion process. We hypothesized that such rare and prolonged encounters arise when an extruder loads approximately midway between the two monomers and draws them progressively into a sphere of radius 𝑅_𝑒_. The encounter persists until continued extrusion extends the loop beyond this range, ultimately displacing the monomers from the encounter sphere (**Fig. 1H**). In this scenario, extrusion reduces the effective genomic separation between loci separated by a linear distance 𝑠 to a smaller value 𝑠’ < 𝑠 (**Fig. 1H**, left). When their average spatial distance 〈(𝑠’)〉 becomes small enough (i.e. 〈𝑅(𝑠’)〉 ∼ < 𝑅_𝑒_) the two monomers are within the encounter radius virtually all the time (**Fig. 1I**). The encounter begins when the effective separation reaches a critical value 𝑠^∗^, defined by the condition 〈(𝑠^∗^)〉 ≈ 𝑅_𝑒_, and terminates once the loop extends by an additional 𝑠^∗^ in the opposite direction. Accordingly, the duration of these loop-extrusion mediated encounters is determined by the time required for the extruder to proceed approximately by 2𝑠^∗^. Since waiting time between consecutive extrusion steps follow an exponential distribution, the time required for 2𝑠^∗^ steps is described by an Erlang distribution (*37*) with a mean time of 2𝑠^∗^/𝑣 where 𝑣 is the loop extrusion velocity. Strikingly, this theoretical prediction accurately predicts the observed distribution of long-lived encounters in simulations where extruders are allowed to move past each other (**Fig. 1J**), as well as the dependence of encounter durations on both 𝑅_𝑒_ and 𝑣 (**Supplementary Fig. 1D-E**). Collisions between extruders in turn lead to a pronounced increase in long encounter durations, reflecting the effective slowdown of extrusion which is not accounted for in this simple theoretical argument (**Fig. 1J**).

In the absence of loop extrusion, the distribution of encounter durations remained largely invariant with respect to linear separation (**Supplementary Fig. 1F**). As a result, when comparing loci at different separation but equal average physical distances, long-lived contacts are observed exclusively in the presence of extrusion (**Fig. 1K)**, further underscoring that these extended interactions reflect an active extrusion-driven mechanism rather than equilibrium polymer dynamics.

We note that while the absolute values of encounter durations scale with the chosen encounter radius - with larger radii yielding longer average contact times - the distributions with and without loop extrusion remain clearly distinguishable across all radii (**Supplementary Fig. 1G**). Importantly, these results are not specific to the ring polymer model we used in the simulations. The same qualitative behavior was observed in simulations using an open-chain Rouse model (**Supplementary Fig. 1H**), indicating that the observed effects reflect general principles of polymer dynamics rather than model-dependent features.

Collectively, these results indicate that the vast majority of chromosomal encounters, even in the presence of loop extrusion, are highly transient and arise from stochastic polymer fluctuations rather than active extrusion by cohesin. In contrast, extrusion-associated encounters are rare, occurring only when an extruder is loaded and initiates extrusion approximately midway within the genomic interval between two loci (**Fig. 1H**). However, these events are markedly longer, and their frequency and duration are sensitive to both the encounter radius and the extrusion velocity. Unlike the waiting times between encounters - which primarily reflect polymer compaction - the distribution of encounter durations thus constitutes a distinct and quantifiable signature of active loop extrusion.

### Defining experimental conditions allowing to observe the dynamic signature of loop extrusion

We next asked whether the long-lived encounters predicted by our model could be experimentally detected in living cells. Validating this prediction would require time-resolved measurements of the 3D distance between two genomic loci, both in the presence and absence of loop extrusion, followed by quantification of encounter durations - defined as consecutive timeframes during which the loci remain within a predefined proximity threshold. However, such measurements do not allow to recover absolute encounter durations. First, the vast majority of encounters predicted by the model are extremely short-lived - most lasting less than a few seconds - and are barely within reach of the temporal resolutions achievable to date (*24*, *38*). Second, and more importantly, localization error in live-cell microscopy systematically inflates observed distances relative to true spatial positions, and leads to fragmentation of continuous encounters into smaller fragments and creation of new artifactual events, even when the error is smaller than the chosen threshold (**Fig. 2A**; see Ref. (*26*)). For these reasons, estimates of encounter events based on hard thresholds onto experimental distance trajectories (*24*, *39*) are severely affected by these artefacts.

Nonetheless, we reasoned that applying hard distance thresholds - while insufficient to recover absolute encounter durations - could still reveal the dynamic signature of loop extrusion under appropriate experimental conditions. As a first step, we reasoned that projecting 3D inter-locus distances onto the xy plane would reduce experimental noise, since axial (z) localization error represents the largest source of uncertainty in live-cell fluorescence microscopy (*40*). 2D projection, however, introduces apparent proximity events (‘phantom’ encounters) due to the loss of spatial information in the z direction, which lead to an overestimation of the duration of random contacts as observed in polymer simulations (**Fig. 2B**, orange lines). Nonetheless, extrusion-driven encounters are substantially longer than random, transient encounters even in the presence of phantom events; so that the extended tail in the distribution of encounter durations in the presence of loop extrusion remains clearly distinguishable from the non-extruding case even after projection (**Fig. 2B**), irrespective of the linear separation between monomers (**Suppl. Fig. 2A**). Therefore, although projection reduces dimensional accuracy, the improved lateral resolution in the xy plane provides a practical advantage for identifying the dynamic signature of loop extrusion.

We next asked whether the differences in encounter duration distributions that could still be observed after projection into 2D would remain detectable in the presence of experimental error on inter-locus distances. To address this, we introduced different amounts of localization noise in the xy plane in polymer simulations and found that, even under these conditions, the distinction between the loop extrusion and non-extrusion cases remained evident provided that the error on 2D distances was smaller than approximately 50% of the threshold (**Fig. 2C** and **Suppl. Fig. 2B**). Although times spent within a threshold distance remain correlated with the underlying absolute encounter durations, encounters identified following xy projection and in the presence of experimental noise should not be interpreted as true physical contact times. Accordingly, we refer to these as ‘apparent encounters’ throughout the remainder of the study.

Together, these findings suggest that detecting the dynamic signature of loop extrusion is, in principle, experimentally feasible, but might require live-cell imaging of chromosomal loci at extremely high spatial and temporal resolution. Since average distances between pairs of loci within TADs are typically in the range of hundreds of nm (*41*, *42*), encounter thresholds should be substantially smaller (in the range of 100 nm or lower). Thus, 2D distance error should be extremely low, in the range of 50 nm or lower based on the above criterion. To test the feasibility of such measurements, we turned to a mESC line which we previously established to visualize the dynamics of two genomic sites with minimal structural or regulatory confounding effects (*21*). These cells harbor two orthogonal operator arrays (∼140x TetO and ∼120x LacO) separated by 150 kb within the ‘neutral’ *Npr3* TAD on chromosome 15, which lacks active or repressive chromatin states and from which we further removed internal CTCF binding sites (*15*). The two arrays can be visualized as sub-diffraction fluorescent foci upon binding of TetR-tdTomato and a low-affinity LacI variant fused to eGFP (LacI**-eGFP) (*43*) (**Fig. 2D**). In addition, a C-terminal HaloTag-FKBP fusion at the endogenous *Rad21* locus enables inducible degradation of RAD21 upon treatment with dTAG-13 (dTAG) (*44*), thus allowing to measure LacO-TetO distances in live cells in the presence or absence of cohesin.

We previously imaged these cells in 3D using oblique illumination microscopy at a 30-second frame rate, achieving an average uncertainty on LacO-TetO distance of approximately 90 nm in 2D after projection in the xy plane (*21*). Under such conditions, the predicted difference in encounter time distributions would not be detectable using thresholds in the ∼<100 nm range required by the average physical distance between the LacO and TetO (approximately 200 nm after xy projection (*21*)) (**Supplementary Fig. 2B, red square**). To overcome this limitation, we optimized the imaging setup by implementing dual back-illuminated scientific CMOS cameras with 95% quantum efficiency, along with custom bandpass filters, dichroic mirrors, and excitation laser lines tailored to the specific emission spectra of eGFP and tdTomato (see **Methods**). This greatly improved photon budget compared to our previous experiments (*21*) thus enabling rapid acquisition of 3D image stacks spanning 9 z-planes (300 nm spacing) every 2 seconds over a 15-minute interval with minimal photobleaching. This represents a 15-fold increase in temporal resolution compared to our previous setup. Combined with higher signal-to-noise ratios and improved spot detection and localization methods requiring minimal post-processing (see **Methods**), these upgrades allowed for substantially denser sampling of LacO-TetO distances (**Fig. 2E**).

To quantify error on distance measurements in these experimental conditions, we first measured frame-to-frame differences in 2D (xy) distances. Mean 2D displacements were on average of 47 nm and 61 nm in the presence and absence of RAD21, respectively, after correction of chromatic aberrations (**Methods**) (**Fig. 2F**). Both values were substantially higher than the 22 nm measured in fixed cells, and remained stable throughout the 15-minute imaging window, indicating consistent localization precision over the course of each movie (**Fig. 2F**). Thus, despite an average measurement error on 2D distances of 22 nm, displacements measured at 2-second frame rate reflected substantial amounts of underlying chromatin dynamics and its changes upon depletion of RAD21. To provide an additional independent estimate of experimental error, we then generated a control cell line containing a single ∼140x TetO array bound simultaneously by TetR-eGFP and TetR-tdTomato (**Fig. 2G**). In this configuration, the green and red signals are expected to co-localize, although heterogeneity in repressor binding across the array may introduce minor separations. Spot detection and localization followed by distance measurement in these control cells yielded a median 2D distance between channels of 37 nm (**Fig. 2H**). We concluded that our experimental error on 2D distances was within 22 and 37 nm, about 2.5- to 4-fold smaller than we could previously achieve (*21*), and in the same range of recent reports (*24*, *38*). Thus, based on the criterion we identified above (**Fig. 2C**, **Suppl. Fig. 2B**), distributions of apparent encounter durations with and without loop extrusion should remain readily distinguishable provided that the (2D) encounter radius exceeds ∼80 nm, i.e. twice the most conservative estimate of error on distances (37 nm).

### Improved live-cell fluorescence microscopy allows experimental validation of model predictions

To test these predictions, we analyzed 786 and 632 LacO-TetO trajectories in untreated cells and cells treated with dTAG-13, respectively, representing 301’695 and 241’574 inter-locus distances acquired across 123 and 213 15-minute movies, respectively. To isolate the dynamic contribution of loop-extruding cohesin and minimize potential confounding from cohesive cohesin complexes, we restricted our analysis to *bona fide* G1-phase cells defined as those without doublet signals indicative of replicated alleles (see **Methods**). As expected, mean projected xy distances between the arrays increased following cohesin loss, rising from approximately 170 nm to 270 nm (**Supplementary Fig. 3A**). Consistent with previous findings (*21*, *22*, *45*), we also observed an increase in the mean squared displacement of the inter-operator distance upon RAD21 depletion, reflecting increased chromatin mobility (**Supplementary Fig. 3B**).

Given the constraints on experimental error and LacO-TetO distances, we then defined a series of threshold distances between 80 and 140 nm and computed the distributions of apparent encounter durations after projecting the trajectories onto the xy plane. In line with model predictions, the presence of RAD21 was associated with a marked increase in the number of long-duration events (**Fig. 3B**), with apparent encounter times extending up to twice the maximum observed under RAD21-depleted conditions regardless of the specific cutoff radius used (**Supplementary Fig. 3C**). However, we note that this difference cannot be unambiguously attributed to loop extrusion: the slower chromatin dynamics observed in the presence of RAD21 may also increase the apparent duration of thermally driven (non-extrusion-mediated) events. These could inflate the tail of the distribution, thereby confounding direct interpretation of the effect of loop extrusion.

**Figure 3:**
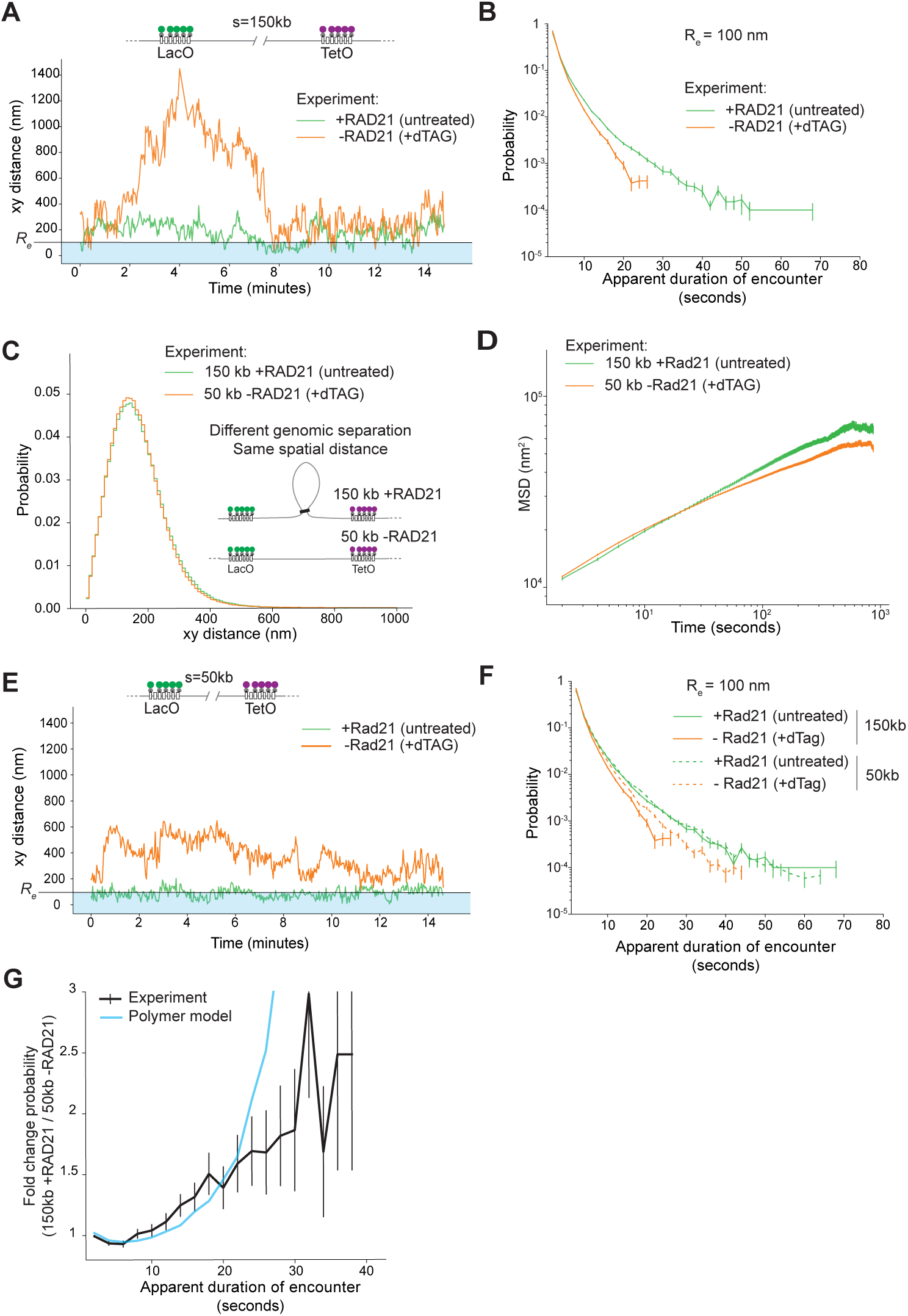
Experimental detection of the dynamical signature of loop extrusion. **A.** Representative experimental trajectories of 2D distances between LacO and TetO arrays separated by 150kb, before (untreated) and following (+dTAG) 2 hours of degradation of RAD21. **B.** Distributions of apparent encounter durations (with 𝑅_𝑒_ = 100 nm) revealing a marked increase in long-lived events in the presence of RAD21. **C.** Overlapping distributions of 2D LacO-TetO distances in untreated cells with an intervening separation of 150kb, and in a derived cell line with 50kb separation, but following RAD21 depletion. **D.** Radial mean squared displacement (rMSD, calculated as the MSD of the vector distance) for the two configurations shown in panel c, demonstrating comparable chromatin dynamics. **E.** Example trajectories from the cell line with 50-kb separation in untreated and RAD21-depleted cells. **F.** Distributions of apparent encounter durations for all combinations of genomic separation (150 kb and 50 kb) and RAD21 depletion status, showing prolonged encounters specifically in the presence of RAD21. **G.** Fold-change in apparent encounter durations between the 150-kb +RAD21 and 50-kb -RAD21 conditions compared to model predictions.

To better control for differences in chromatin mobility, we generated a new mESC line in which the separation between the LacO and TetO arrays was reduced to 50 kb by deleting a 100-kb segment of intervening ‘neutral’ genomic sequence. In this modified line, both the mean physical distance between the arrays and the mean squared displacement (MSD) in the absence of RAD21 closely matched those of the original 150-kb system in the presence of RAD21 (**Fig. 3C-D**). Despite this equivalence in spatial and dynamic properties, the distributions of apparent encounter durations remained distinct: in the absence of RAD21, encounters were significantly shorter than those observed in the original cell line with RAD21. In the presence of RAD21 however, the distributions were nearly identical across both cell lines (**Fig. 3F**, **Suppl. Figure 3D**). These results indicate that the enrichment of long-lived apparent encounters observed in the presence of RAD21 is not due to slower chromatin dynamics, but rather specifically reflects the association between the base of cohesin-held loops. Supporting this interpretation, the experimentally observed fold-change in apparent encounter duration between the untreated 150-kb configuration and the RAD21-depleted 50-kb condition closely matched model predictions (**Fig. 3G, Suppl. Figure 3E**).

Thus, although measurement noise and 2D projection preclude direct access to true encounter durations, the substantially improved spatial and temporal resolution of our imaging approach enabled us to validate key predictions from polymer simulations and provided experimental evidence of long encounter times that are the dynamic signatures of loop extrusion.

### A model of loop-extrusion mediated transcriptional activation

Taken together, our theoretical and experimental findings demonstrate that the vast majority of encounters between genomic loci are driven by stochastic fluctuations of the chromatin fiber and are extremely short-lived. In contrast, a small subset of encounters is dependent on loop extrusion and exhibits significantly longer durations, extending up to several minutes and dependent on the encounter radius (**Suppl. Fig. 1G**) and extrusion velocity (**Suppl. Fig. 1E**). This non-trivial distribution of encounter durations prompted us to ask whether such dynamics might influence enhancer-promoter communication.

If all enhancer-promoter encounters were equally ‘productive’ (that is, capable of eliciting a transcriptional response regardless of their duration) then transcriptional output over a given time interval would be expected to scale linearly with the overall encounter probability between the enhancer and promoter. However, this is inconsistent with previous findings from our lab and others (*10*, *14–16*), which demonstrated that transcriptional outputs exhibit nonlinear relationships with enhancer–promoter contact probabilities. These nonlinearities have previously been attributed to molecular mechanisms in which frequent and short-lived enhancer-promoter encounters are integrated through a sequence of rate-limiting regulatory steps (*15*), or through promoter ‘futile cycles’ that reduce the efficiency of activation despite frequent contacts (*14*).

An alternative and opposite explanation, also recently proposed by another study (*46*), is that only encounters exceeding a minimum duration threshold are transcriptionally productive (**Fig. 4A**). Such a requirement could arise from kinetic proofreading (*47*) or rate-limiting assembly steps at the promoter (*48*) requiring the simultaneous presence of multiple transcription factors or cofactors at the enhancer-promoter interface (*49*, *50*). A similar outcome could also arise if long-lived transcription factor binding events are required for efficient enhancer-promoter communication and to initiate transcription (*51*). In both cases, the likelihood of observing productive transcription would naturally increase with encounter duration. We therefore set out to explore whether transcriptional events mediated by the rare and long-lived encounters mediated by loop extrusion could explain the experimentally observed nonlinearities, and how transcription responds to depletion of cohesin subunits or associated cofactors (*10–13*).

**Figure 4:**
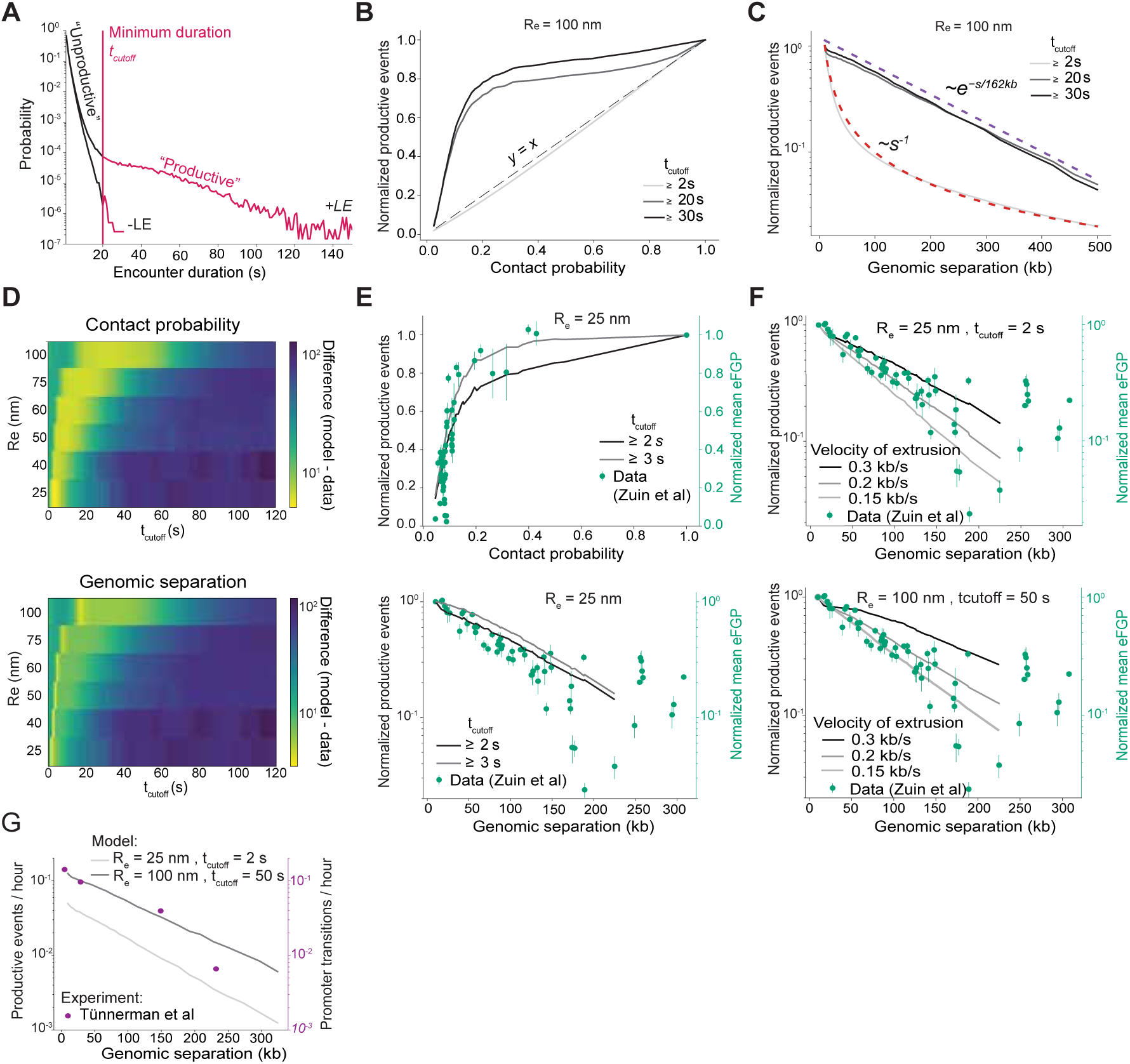
A minimal model of transcriptional activation by rare, long-lived, enhancer-promoter encounters mediated by loop extrusion. **A.** Schematic of the model where only enhancer-promoter encounters exceeding a minimum duration threshold (𝑡_𝑐𝑢𝑡𝑜𝑓𝑓_) may be transcriptionally productive. **B.** Polymer simulations with loop extrusion reveal that increasing the threshold for productive encounter duration progressively introduces a nonlinear relationship between the frequency of productive encounters and overall contact probability. **C.** The scaling behavior of productive encounters shifts from a power-law, red dotted line, to an exponential decay, purple dotted line, with genomic distance as 𝑡_𝑐𝑢𝑡𝑜𝑓𝑓_ increases. **D.** Heat map quantifying agreement between model predictions and experimental transcriptional output measured in Ref. (*15*) as a function of contact probability (top panel) and as a function of genomic separation (lower panel), for a range of encounter radii and values. Absolute best agreement is achieved for a 25 nm radius and 2 to 3 s durations. **E.** Productive encounters, for 𝑅_𝑒_ = 25 nm and 𝑡_𝑐𝑢𝑡𝑜𝑓𝑓_ = 2 and 3 s, plotted as a function of contact probability (top panel) and genomic separation (bottom panel) recapitulate respectively the observed non-linear relationship between transcriptional output and contact probability and the exponential decay of transcriptional output with genomic distance (experimental data from Red. (*15*)). **F.** Productive encounters as a function of genomic separation, predicted by the model for 𝑅_𝑒_ = 25 nm and 𝑡_𝑐𝑢𝑡𝑜𝑓𝑓_ = 2 s (top panel) and for 𝑅_𝑒_ = 100 nm and 𝑡_𝑐𝑢𝑡𝑜𝑓𝑓_ = 50 s, when varying the velocity of extrusion. **G.** Predicted frequency of productive encounters as a function of genomic distance for 25 nm radius, 2 s duration and 100 nm radius, 50 s duration from the polymer model with extrusion velocity equal to 0.2 kb/s. Data points: *Sox2* promoter transition rates from low-frequency to high-frequency bursting states inferred from prior live-cell imaging of transcriptional dynamics (*19*).

To investigate this scenario, we reanalyzed our polymer simulations with loop extrusion by plotting the number of 3D encounters exceeding arbitrary minimum durations against the overall contact probability, for pairs of monomers separated by varying genomic distances. As expected, when the minimum duration was low (effectively treating all contacts as potentially productive) the relationship was linear. However, increasing the threshold for productive encounter duration progressively resulted in nonlinear behaviors closely resembling those observed experimentally (*15*) (**Fig. 4B**). This behavior, which occurs for any choice of the value of the encounter radius, is a direct consequence of increased enrichment for loop-extrusion mediated encounters. While the overall contact probability between loci scales as a power law with genomic distance (*52*) (**Fig. 1A**), the probability that two loci are bridged by an active extruder instead decreases exponentially (**Supplementary Fig. 4A**). The rate of this exponential decay is equal to the extruder processivity (speed x lifetime) in the absence of collisions (**Supplementary Fig. 4A**, orange line) whereas it is lower in the presence of collisions due to the effective lower processivity of the extruder (**Supplementary Fig. 4A**, blue line). In both cases, this effect decouples the probability of long productive transcriptional interactions from the total contact probability, producing a nonlinear relationship as seen in **Fig. 4B**. Importantly, this framework also predicts that as the minimum duration required for a productive encounter increases, the scaling of the frequency of productive events with genomic distance should shift from a power-law to an exponential decay (**Fig. 4C**). Consistent with the interpretation that this results from loop-extrusion driven encounters, the decay rate of the exponential is slightly lower to the processivity of the extruder (162 kb vs. a processivity of 180 kb in **Fig. 4C**).

We next asked whether this minimal model could quantitatively recapitulate experimental observations. Using polymer simulations parameterized to reproduce contact probabilities in the presence of RAD21 (**Fig. 1A**), we systematically varied the encounter radius and the minimum duration required for an encounter to be considered productive (ranging from 25 to 100 nm and 0 to 120 seconds, respectively). We then compared the predicted nonlinear relationships to our previously published measurements of transcriptional output from the *Sox2* promoter, in which the SCR enhancer was relocated to multiple genomic positions within the same neutral TAD (*15*). The model closely reproduced the experimental data across a broad parameter space, with the best agreement achieved for an encounter radius of 25 nm and minimum durations between 2 and 3 seconds (**Fig. 4D**). Similar levels of agreement with the experimental data were obtained across a range of encounter radii, each associated with a distinct range of optimal minimum durations that increased with increased encounter radius (**Fig. 4D**). Strikingly, we also observed that transcriptional output from the *Sox2* promoter decreased exponentially with increasing genomic distance from the enhancer, closely matching model predictions across the same range of encounter radii and minimum encounter durations (**Fig. 4E**). Quantitative agreement with experimental data was further improved by a moderate reduction in the extrusion velocity used in the simulations (**Fig. 4F**). Together, these findings suggest that, regardless of the spatial scale at which functional enhancer–promoter communication occurs, rare but sustained loop-extrusion–mediated encounters are sufficient to explain the transcriptional response of the *Sox2* promoter as a function of enhancer-promoter genomic separation.

As expected, for all parameter combinations that recapitulate the experimental data, the minimum duration required for a productive encounter consistently exceeds that of the vast majority of random contacts and mostly captures loop extrusion-mediated events. This is further supported by the observation that the optimal minimum duration falls within the same range as the typical time required to extrude a pair of monomers across the corresponding encounter radius (2𝑠^∗^/𝑣 ≈ 3 to 40 seconds) (**Supplementary Fig. 4B**). Therefore, an additional prediction of this model is that productive encounters between an enhancer and a promoter should be extremely rare, regardless of the spatial scale at which enhancer–promoter communication occurs. For the parameter set corresponding to the absolute best fit of experimental data (25nm radius and 2s minimum time), predicted encounter frequencies ranged from approximately 0.05 to 0.001 events per hour at 10 and 300 kb separation, respectively, and increased to 0.1 and 0.01 per hour, respectively, in case a an encounter radius of 100 nm was considered with 50 s minimum time (**Fig. 4G**). Notably for loci separated by 150 kb as in the live-cell imaging experiments reported in **Fig. 2-3**, productive encounters within a radius of 100 nm should occur only 0.04 times per hour - corresponding to slightly less than one event per 16-hour cell cycle in mESCs. Testing this prediction would require simultaneous live imaging of enhancer and promoter positions along with transcriptional output (or related molecular signatures) at the promoter, under conditions where experimental error is minimized to a level that allows unambiguous detection of absolute encounter durations. At present, such measurements remain unachievable for the reasons exposed above. Interestingly however, these predicted low frequencies of productive encounters resembles those at which we recently inferred the *Sox2* promoter to switch from a basal to a frequently-bursting regime, based on live-cell imaging of transcriptional dynamics as a function of enhancer-promoter separation (*19*) (experimental points in **Fig. 4G**).

Taken together, these results indicate that an enhancer’s regulatory influence on its target promoter may be exerted only infrequently during the cell cycle, primarily through rare, transiently sustained encounters driven by loop extrusion.

### Prediction of transcriptional effects of depleting cohesin and cofactors

We next tested the ability of this simple model to predict the transcriptional consequences of experimental perturbations to the dosage of cohesin or its cofactors, which alter its DNA occupancy or loop extrusion dynamics. We specifically focused on predicting the effects of depleting cohesin and NIPBL, for which the impact on promoter activity has been experimentally quantified as a function of genomic distance from an enhancer (*10–13*). To simulate cohesin depletion, we decreased the density of loop extruders progressively, and observed that this increasingly disrupted the exponential decay in the frequency of productive encounters as a function of genomic separation (**Fig. 5A**). In conditions that mimic full depletion conditions (no extrusion), the model predicted that productive encounters (and by extension, transcriptional activity) should be rarer and decay not as an exponential but rather as a power law, mirroring the power-law decay of overall contact probabilities (**Fig. 5A**). This is because in the absence of loop extrusion, transcription uniquely relies on the occasional occurrence of random encounters whose duration exceeds the minimum required, and the probability of such events for any combinations of encounter radius and minimum duration is proportional to overall contact probabilities. This prediction is consistent with recent observations in *Drosophila*, where chromosomes are likely not folded by the loop extrusion activity of cohesin, and where burst frequency at the *Even skipped (eve)* gene was shown to decrease as an inverse power-law of its genomic separation with an ectopic copy of its enhancer (*53*). The model also predicted that even partial depletion of extruder dosage should result in reduced transcriptional outputs, with the magnitude of this effect growing rapidly and then saturating to a constant value as the genomic distance between enhancer and promoter grows (**Fig. 5B**). In contrast, the model predicted little to no change in transcriptional output following cohesin depletion when the enhancer-promoter separation is sufficiently short such that the loci remain within the encounter radius at all times (that is, approximately ∼2 kb for a 25 nm radius, or 5 kb for a 100 nm radius (**Suppl. Fig.5A**)). These predictions are again consistent with experimental results from inducible depletion of cohesin (*10–12*, *54*).

**Figure 5:**
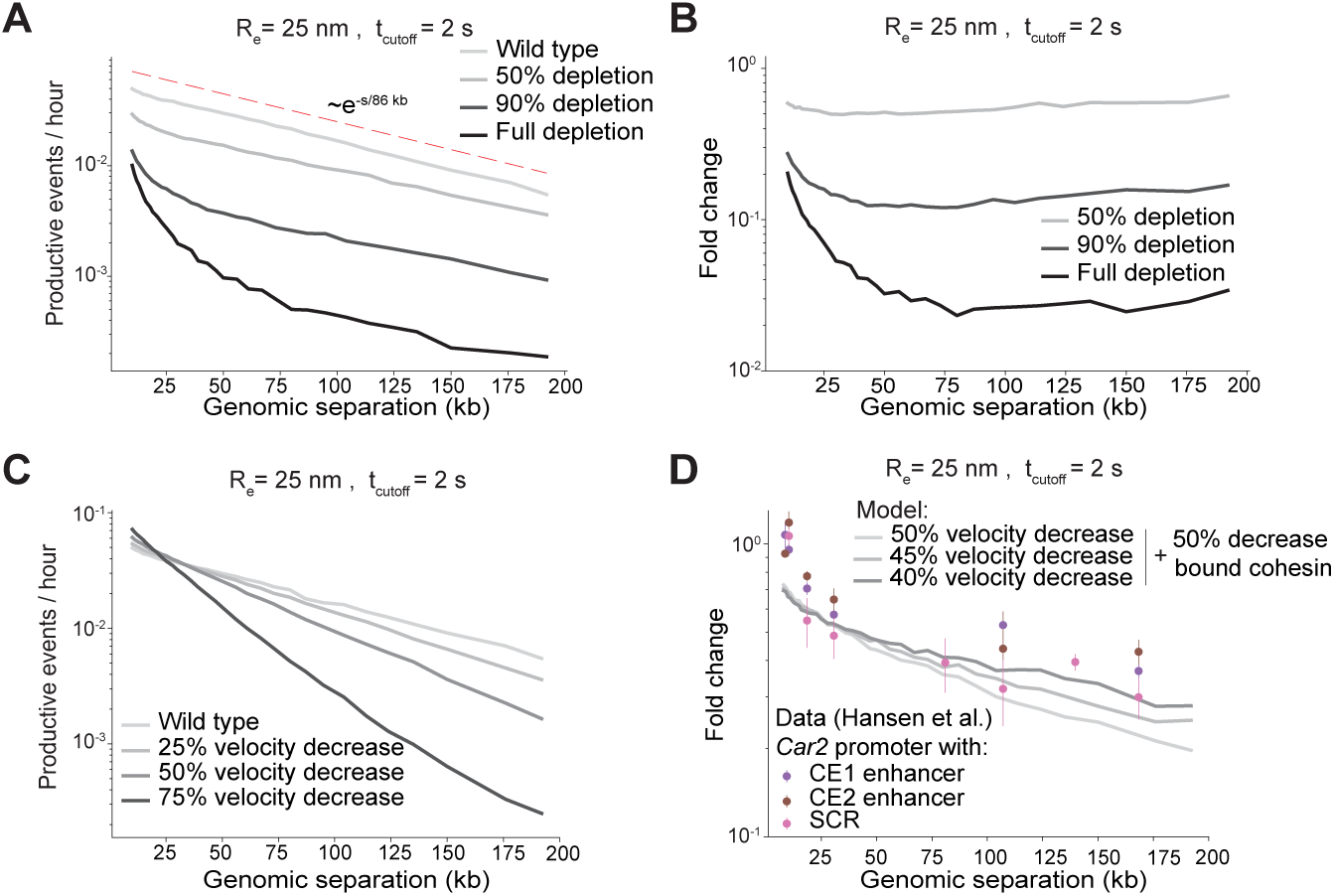
Prediction of the transcriptional effects of cohesin and NIPBL depletion. **A.** Simulated distributions of productive enhancer–promoter encounter frequencies as a function of genomic distance when varying loop extruder density to simulate cohesin depletion. Progressive depletion of extruders (light to dark gray) results in productive encounters switching from exponential to power-law decay as a function of genomic separation (without extrusion). Here productive encounters are defined as those lasting more than 2s at a radius of 25 nm. **B.** Predicted fold-change in transcriptional output following partial cohesin depletions across increasing enhancer-promoter distances, with same parameters as in panel A **C.** Simulated effect of decreasing extrusion velocity on productive encounter frequency. Slower extrusion increases encounter probability at short genomic distances but reduces it at longer distances, while maintaining the exponential form of distance-dependent decay until 90% decrease. Same parameters as in panel A. **D.** Transcriptional fold changes predicted by the model following NIPBL depletion, incorporating experimentally measured reductions in both cohesin density and velocity (*55*). The model was set to 50 % decrease in bound cohesin and varying extrusion velocity form (50%, light gray to 37.5% in dark gray). Experimental data: previously reported fold change in transcriptional outputs between cells treated with dTAG-13 for NIPBL depletion and untreated cells from the *Car2* promoter as a function of its genomic separation from either of its endogenous enhancers (CE1 and CE2) or the Sox2 control region (SCR) enhancer. Parameters as in panel A.

We next simulated the effects of NIPBL depletion, which has been shown to reduce both cohesin loading and extrusion velocity in vivo (*55*). In the model, decreasing extrusion velocity led to a modest increase in the number of productive encounters at short genomic distances and a pronounced reduction at longer distances (**Fig. 5C**). This dual effect arises because slower extrusion rates, with constant extruder residence time, yield smaller loops; and while shorter loops enhance the probability of productive encounters at short range, they diminish the likelihood of extrusion-mediated contacts over extended distances. Notably, unlike the loss of loop extruders, reduced extrusion velocity preserved the exponential decay of transcriptional output with increasing enhancer–promoter separation (**Fig. 5C**). Strikingly, when we simulated the experimentally measured ∼50% reductions in both cohesin density and extrusion speed observed in NIPBL-depleted mESCs treated with dTAG-13 (*55*), the model predicted the transcriptional fold changes at the *Car2* promoter as a function of its genomic distance to either of its endogenous enhancers or to the Sox2 Control Region (SCR), as recently reported in ref. (*13*), without any parameter fitting (**Fig. 5D**). The model notably predicted that in contrast with cohesin depletion alone, the transcriptional effect of NIPBL depletion grows with increasing genomic distance. Together, these results further support the notion that rare, but sustained encounters between enhancers and promoters mediated by the loop extrusion activity of cohesin drive their functional communication.

## Discussion

Our study provides theoretical and experimental evidence for a previously uncharacterized class of encounters between genomic loci that specifically depend on the loop extrusion activity of the cohesin complex. These encounters occur when cohesin is loaded approximately midway between two genomic regions, subsequently extruding them through a defined encounter region. Although considerable uncertainty exists regarding the spatial proximity required for an enhancer to activate transcription at a promoter (ranging from tens to hundreds of nanometers depending on the underlying regulatory mechanisms (*56*, *57*)) our results show that extrusion-driven encounters consistently remain markedly longer-lived than random chromatin fiber collisions across this entire range. With an extrusion velocity of 0.3 kb/s, within the expected range in living cells (*21*, *22*), and encounter radii spanning 25 to 200 nm, extrusion-mediated encounters are predicted to last between 3 and 130 seconds (**Supplementary Fig. 4B and Supplementary Fig.1D**). While substantially longer than random collisions, which typically last 1 to 4 seconds in the same range of radii, such encounters remain an order of magnitude shorter than CTCF-anchored configurations of the chromatin fiber, which we and others previously showed to last approximately 10 to 30 minutes (*21–23*).

The spatial and temporal scales of encounters due to loop extrusion closely match the limits of frame rates and localization precision achievable in dual-color imaging of chromosome dynamics, which inherently prevents direct measurement of absolute encounter durations (**Fig. 2A**) and confounds previous estimations including from our own lab (notably the ‘proximal’ state detected without CTCF sites in Ref. (*21*)). Nevertheless, we identify experimental conditions under which exceptionally high spatial and temporal resolution, combined with two-dimensional projection of motion, allow for the detection of a clear signature of extrusion-driven encounters. This signature is notably absent in cells lacking cohesin. Although loop extrusion also shortens the time between successive encounters, as recently observed in live-cell studies (*24*, *25*), our analysis indicates that this effect arises solely from increased chromatin compaction resulting from cohesin-mediated bridging of distal loci at the bases of extruded loops, rather than from the active extrusion process itself. The rare and sustained physical encounters revealed in our study therefore represent the only direct dynamic hallmark of loop extrusion.

Our findings have important implications for transcriptional regulation by distal enhancers. The simple assumption that regardless of the exact encounter radius, longer-lived (and thus, mostly extrusion-driven) encounters are more likely to lead to productive transcription provides a unifying model for understanding how contact probabilities are nonlinearly translated into transcriptional outputs, as recently observed in multiple studies (*10*, *14–16*). It also allows to rationalize how changes in cohesin concentration or in the dosage of cofactors that modulate its loop extrusion activity affect transcription levels, particularly in genomic contexts where confounding structural and regulatory influences are minimized (*10–13*). Our findings complement and extend a recent report proposing that time-gated encounters can generate nonlinear transcriptional responses (*46*) by uniquely identifying loop extrusion-mediated encounters as the predominant class of events whose durations match those required to quantitatively recapitulate experimental observations (**Fig. 4-5**).

This model further predicts that an enhancer should exert a regulatory effect on its target promoter only infrequently, ranging from a few events per cell cycle on average to less than one, depending on genomic separation. While this may appear counterintuitive and contrasts with earlier models that attributed enhancer function to frequent, brief random encounters followed by slower regulatory steps (*14*, *15*), it aligns closely to our recent live-cell measurements of *Sox2* promoter bursting dynamics at varying genomic distances from its SCR enhancer (*19*). There, we showed that the presence of the enhancer triggers rare, stochastic transitions in the promoter from extended periods of low-frequency bursting to periods of high-frequency activity. The frequency of these transitions strikingly matches the predicted frequency of loop extrusion-driven encounters for the corresponding genomic separations (**Fig. 4G**). This allows us to identify the source of promoter regime transitions with the long-lived, mainly loop extrusion-driven encounters, which thus would control promoter burst frequency not by modulating the occurrence of single bursts but rather of long-lived periods of high promoter bursting activity. In this emerging scenario, and at least in genomic regions devoid of further regulatory sequences and CTCF sites, enhancer-promoter encounters would be a major rate-limiting step in transcriptional activation, and would set fundamental constraints on whether or not a promoter can be regulated by an enhancer in the course of a single cell cycle.

In this work, we did not consider that enhancer or promoter sequences might partially block cohesin, for example *via* interactions with RNA Pol II (58). This would not affect the conclusions of our work, but we note that in such a scenario loop-extrusion mediated encounters between the enhancer and promoter would become more frequent, because they would occur even when cohesin is loaded further from the midpoint. A limit case of such a scenario is provided by the simulations shown in **Suppl. Fig. 1C,** where impermeable extrusion barriers emulating CTCF sites led to a substantial increase in the frequency of cohesin-driven encounters. We also did not consider the possibility that enhancer-promoter encounters might be stabilized by attractive molecular interactions (*31*, *59*, *60*). However, such a mechanism is unlikely to alter the main conclusions of our study. While attractive interactions would generally increase encounter durations and thus enhance the likelihood of productive transcription, the frequency of such events would scale as an inverse power law with genomic distance. Thus, reproducing the experimentally observed nonlinear transcriptional response would still require the exponential distance-dependence introduced by loop extrusion.

Together, our results provide a unifying framework to understand transcriptional regulation by distal enhancers, and indicate that it relies on time-gated molecular processes whose occurrence is boosted by sustained physical encounters under the loop-extrusion activity of cohesin.

## Acknowledgements

We thank Laurent Gelman and Laure Plantard from the FMI Facility of Advanced Microscopy and Imaging (FAIM) for help with microscope setup; all members of the Giorgetti lab for discussions and feedback on the manuscript; Leonid Mirny, Timothy Földes and Elphège Nora for discussing unpublished data; Elphège Nora, Anders S. Hansen, Geoffrey Fudenberg, Tineke Lenstra, Elzo de Wit, David Brückner, Verena Mutzel and Gergely Tihanyi for comments on the manuscript. Research in the Giorgetti lab is funded by the Novartis Foundation and the Swiss National Science Foundation (grants no. 310030_192642, TMCG-3_213782 and 320030-236070). M.U. was supported by an EMBO Postdoctoral Fellowship (ALTF 1305-2024).

## Data availability

The trajectories from imaging data can be found on Zenodo: https://doi.org/10.5281/zenodo.17185366.

## Code availability

All the code generated for this study can be found at: https://github.com/giorgettilab/loop_extrusion_contact_dynamics (loop extrusion simulation and image analysis).

## Author contributions

M.U. conceived the study. and performed polymer simulations, microscopy and data analysis. N.L. performed microscopy and data analysis. K.L. and J.C. generated cell lines. G.R. performed data analysis.

P.I.K. performed initial simulations. E.M. and G.T. developed code for polymer simulations. L.G. conceived and supervised the study and wrote the paper with M.U. and N.L. and input from all authors.

## Supplementary Materials

One document with:

Materials and Methods

Supplementary Table 1: Oligonucleotides

Supplementary Table 2: Numbers of experimental replicates

Supplementary references

---

**Supplementary Figure 1:**
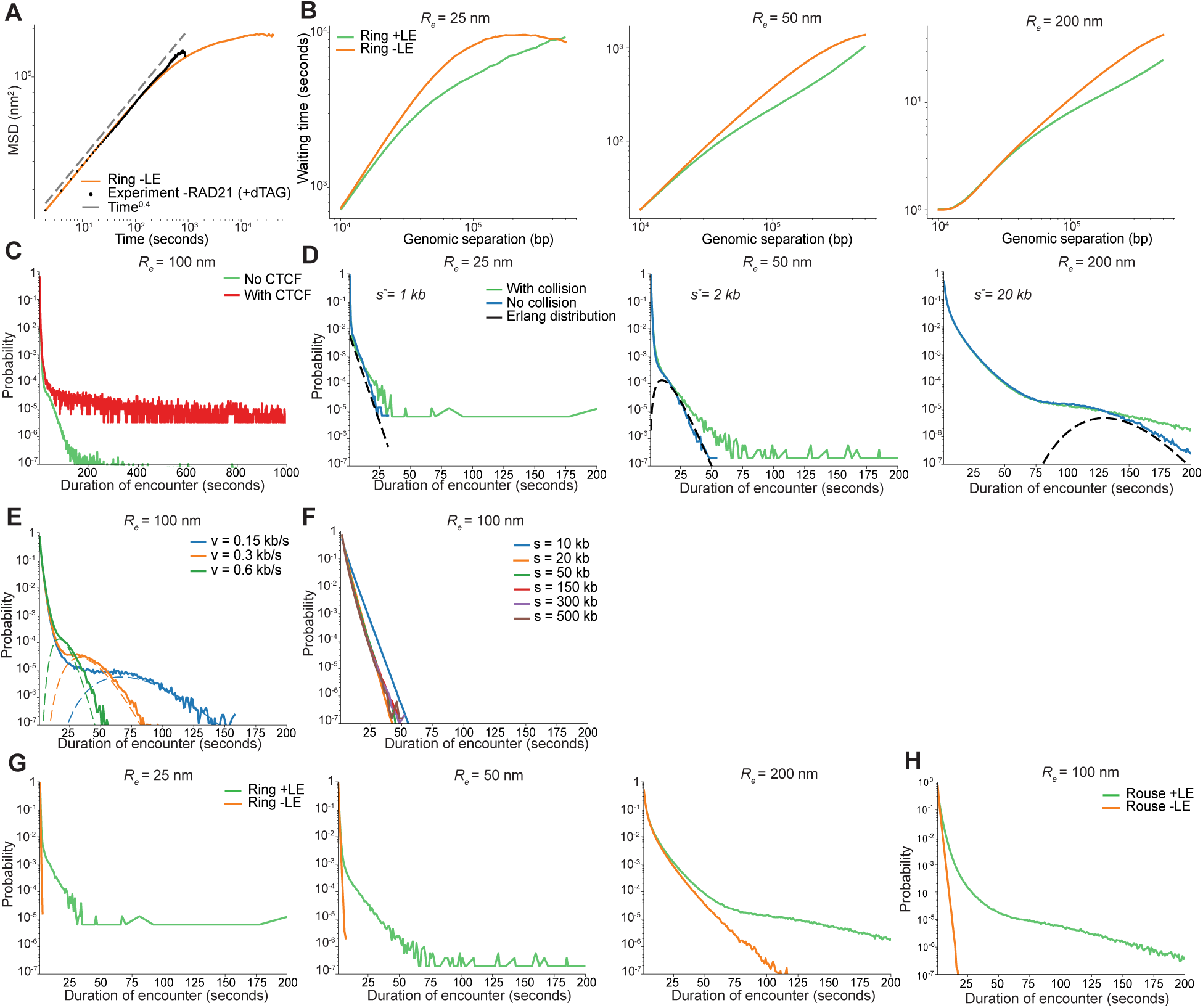
**A.** Mean squared displacement (MSD) of the 2D vector distance between genomic loci separated by 150 kb in the ring polymer model reproduces the subdiffusive behavior observed in live-cell imaging experiments reported in Fig. 2, with a scaling exponent of ∼0.4. Experimental MSD was measured in mESC treated for 2 hours with dTAG-13 for RAD21 depletion and imaged for 15 minutes every 2 seconds in 3D (9 stacks separated by 0.3 µm, 100ms exposure time; see Fig.2, related text and Methods section for details). **B.** Average waiting times between encounters in the ring polymer plotted as a function of genomic distance for various encounter radii 𝑅_𝑒_, with and without loop extrusion. **C.** Distribution of encounter durations in the absence and presence of extrusion barriers. **D.** Distribution of encounter durations for different encounter radii with or without collision of extruders. Black dotted lines: corresponding Erlang distributions with mean values 2𝑠^∗^/𝑣 and rate 1/𝑣 (see Fig. 1H). **E.** Distribution of encounter durations for different extrusion velocities, plotted together with the corresponding Erlang distributions, illustrating that they closely match the long-time behavior of distributions. **F.** Distribution of encounter durations in absence of loop extrusion for different genomic separation. **G.** Comparison of distributions of encounter durations with and without loop extrusion across a range of encounter radii. **H.** Distributions of encounter durations in an open-chain Rouse polymer model with and without loop extrusion.

**Supplementary Figure 2:**
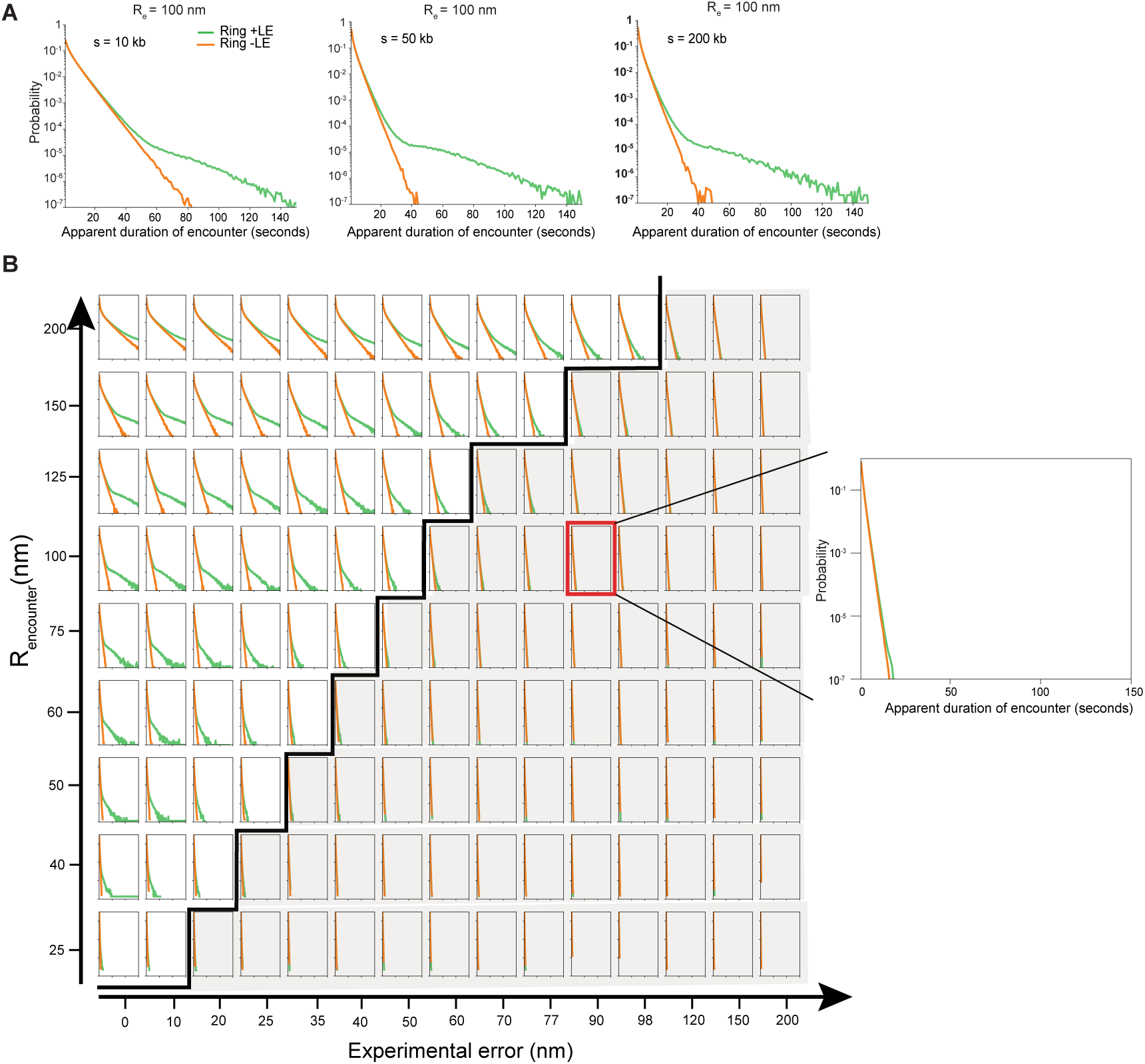
**A.** Simulated distribution of apparent encounter durations for monomers separated by respectively 10, 50 and 200kb on a ring polymer with or without loop extrusion. The encounter radius chosen was 100 nm in 2D. **B**. Distribution of apparent encounter durations for monomers separated by 150 kb on a ring polymer with or without loop extrusion for different encounter radii between monomers ranging from 25 to 200 nm varying the experimental error (0 to 200 nm) on the 2D distance. The error is simulated by applying random gaussian variables on each component of the vector distance. Every subplot corresponds to one combination of encounter radius and experimental error. Black line represents the line below which the distribution with or without extrusion qualitatively differs from one another which corresponds to errors larger than approximately 𝑅_𝑒_/2. The red square, magnified on the right, indicates distribution of encounter duration with a radius of encounter of 100 nm and experimental error of 90 nm as in Mach et al.

**Supplementary Figure 3:**
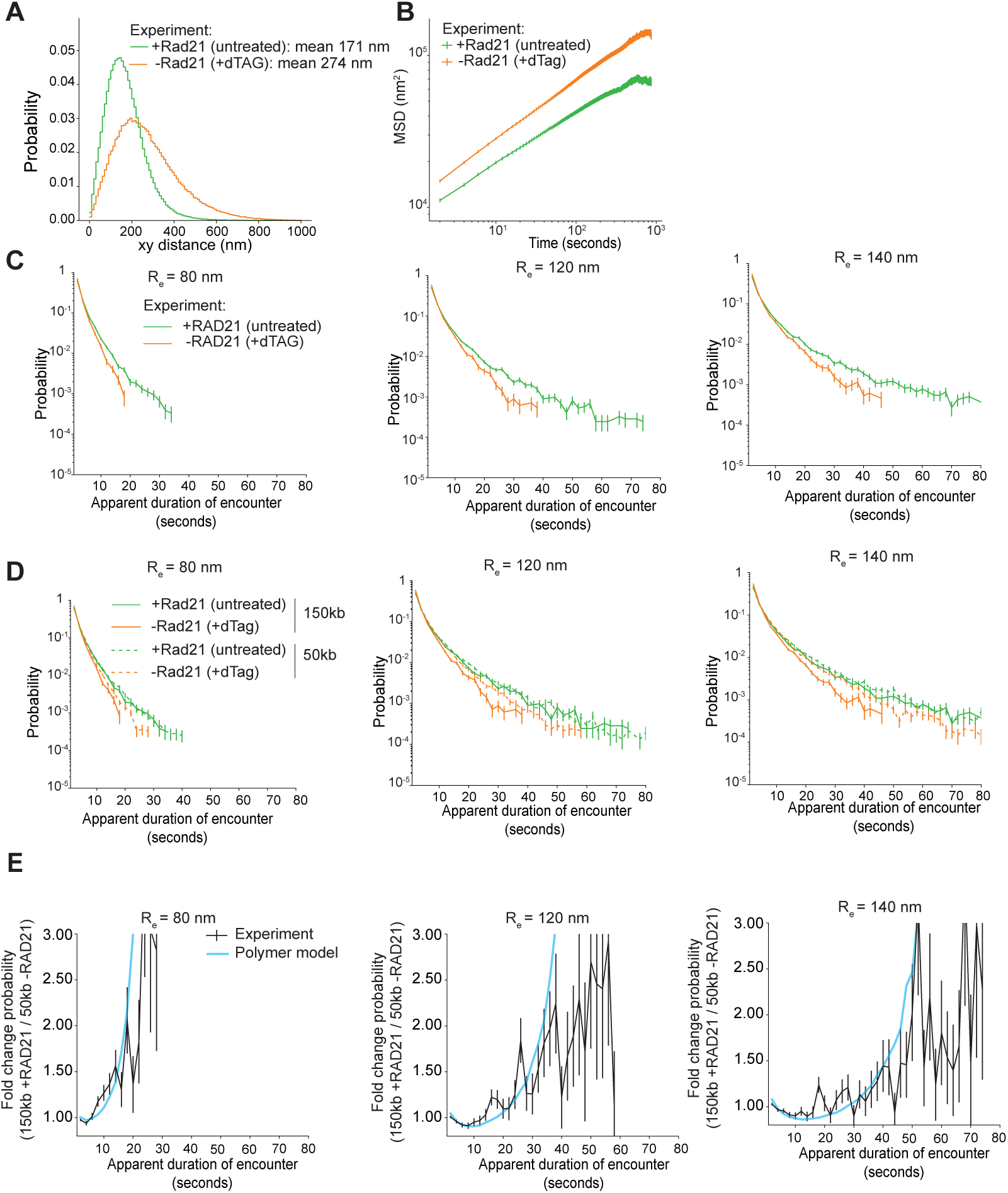
**A.** Distribution of 2D distances between LacO and TetO for the cell line with 150 kb genomic separation before (untreated) and following (+dTAG) 2 hours of degradation of RAD21. The legend includes the average of the distribution taken from 125 and 233 movies, 840 and 729 tracks for untreated and +dTag respectively. **B.** Mean squared displacement of the 2D vector distance for the cell line with a genomic separation of 150 kb before or after depletion of RAD21. **C.** Distribution of encounter duration for the cell line with 150 kb genomic separation with (green) or without (orange) the presence of Rad21 for three different encounter radius namely 80,120 and 140 nm. **D.** Distributions of apparent encounter durations for all combinations of genomic separation (150 kb and 50 kb) and RAD21 depletion status for three different encounter radius 80,120 and 140 nm. **E.** Fold-change in apparent encounter durations between the 150-kb+RAD21 and 50-kb -RAD21 conditions compared to model predictions for the three encounter radius considered in the previous panel.

**Supplementary Figure 4:**
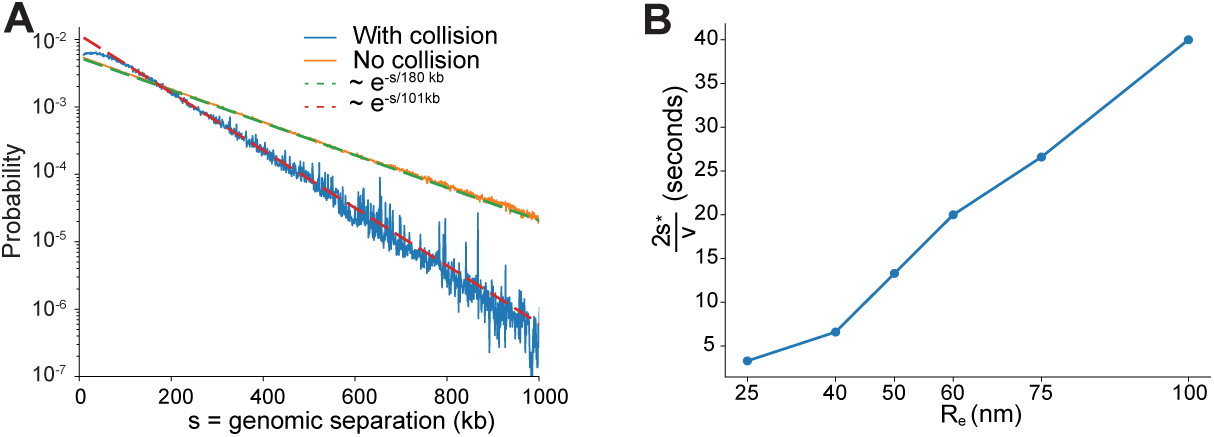
**A**. Probability that two loci are bridged by an active extruder as a function of their genomic separation in the presence (blue) or not (orange) of collisions between extruders. The dotted lines represent the exponential decay of the two curves. In the absence of collision, the distribution decays, as expected, with a rate equal to the processivity of the extruder. In the presence of collisions, the rate becomes smaller because the effective processivity of the extruder is shorter. **B.** Average value, 2𝑠^∗^/𝑣, of the Erlang distribution describing the tail of the distribution of the encounter duration, shown in Figure 1 and **Supplementary Figure 1**, as a function of the encounter radius.

**Supplementary Figure 5:**
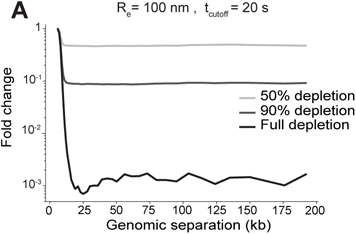
**A.** Predicted fold change in transcriptional output following partial cohesin depletion (from 50 % depletion to full depletion, light to dark gray) across increasing enhancer-promoter distances. The encounter radius is set to 100 nm in 3D and a time cutoff of 20s is applied to define an encounter.

