## Supplementary Material for "Loop extrusion creates rare, long-lived encounters underlying enhancer-promoter communication"

**This PDF file includes:**

Materials and Methods
Supplementary Table 1: Oligonucleotides
Supplementary Table 2: Numbers of experimental replicates
Supplementary References

### Materials and Methods

#### Culture of mouse embryonic stem cells (mESC)

For genome engineering, cells were cultured on gelatin-coated culture plates in Glasgow Minimum Essential Medium (GMEM) (Sigma-Aldrich, G5154) supplemented with 15% foetal calf serum (Eurobio Abcys), 1%L-Glutamine (Thermo Fisher Scientific, 25030024), 1% Sodium Pyruvate MEM (Thermo Fisher Scientific, 11360039), 1% MEM Non-Essential Amino Acids (Thermo Fisher Scientific, 11140035), 100$\mu$M $\beta$-mercaptoethanol (Thermo Fisher Scientific, 31350010), 20U/ml leukemia inhibitory factor (Miltenyi Biotec, premium grade) in 8% $CO_2$ with 2i inhibitors (1$\mu$M MEK inhibitor PDO35901 (Axon, 1408) and 3$\mu$M GSK3 inhibitor CHIR 99021 (Axon, 1386)) at 37°C. For live-cell imaging experiments, $10^6$ cells were seeded on 35-mm glass-bottom dishes (Mattek, P35G1.5-14-C) coated with 2 $\mu g\ ml^{-1}$ laminin (Sigma-Aldrich, L2020) in PBS at 37 °C overnight and cultured for 24 h in Fluorobrite Dulbecco's Modified Eagle Medium (DMEM) (Gibco, A1896701) supplemented with 15% foetal calf serum (Eurobio Abcys), 1% L-Glutamine (Thermo Fisher Scientific, 25030024), 1% Sodium Pyruvate MEM (Thermo Fisher Scientific, 11360039), 1% MEM Non-Essential Amino Acids (Thermo Fisher Scientific, 11140035), 100$\mu$M $\beta$-mercaptoethanol (Thermo Fisher Scientific, 31350010), 20U/ml leukemia inhibitfory factor (Miltenyi Biotec, premium grade) and 2i inhibitors. For depletion of RAD21, the medium was replaced with Fluorobrite medium containing 500 nM dTAG-13 (Sigma-Aldrich, SML2601-1MG) 2 hours before imaging.

#### Generation of mESC lines

All cell lines are based on E14Tg2a mouse embryonic stem cells (karyotype 19, XY, 129/Ola isogenic background; E14 for brevity). The cell line with TetO and LacO separated by 150kb (1A2) was established in previous work (*1*). LacO and TetO insertion coordinates (mm39) were chr15:11567240 and chr15:11717704, respectively. The cell line with reduced distance (50kb) between the operators (1D9) was generated from a modified version of the 1A2 line in which the homologous untargeted chr15 allele was further engineered by removing the entire 'neutral' TAD (sgRNA sequences in **Supplementary Table 1**). To reduce the distance between operators on the targeted allele, two gRNAs were designed using the IDT online designing tool (https://www.idtdna.com/site/order/designtool/index/CRISPR_SEQUENCE) to target two genomic locations separated by 100kb between the LacO and TetO sequences ( sequences in **Supplementary Table 1**). Cells were cultured as described above. 500'000 cells were transfected with Lipofectamine3000 according to the manufacturer's instructions (Thermo Fisher Scientific, L3000008) using 0.5 µg of each gRNA. The media was changed the day after transfection and six days post-transfection the cells were sorted by FACS into single clones in a 96-well plate. At 12 days after sorting, the plates were merged and genomic DNA was extracted on the plate using a lysis buffer (100 mM Tris-HCl pH 8.0, 5 mM EDTA, 0.2% SDS, 50 mM NaCl and 0.05 mg ml−1 proteinase K (Macherey-Nagel, 740506) and 0.05 mg ml−1 RNase A (Thermo Fisher Scientific, EN0531) followed by an incubation at 55°C for 1h and 5 minutes at 94°C. The clones were then checked for deletion of genomic DNA by PCR (sequences of the primers used can be found in **Supplementary Table 1**). Positive clones were further verified after extraction and purification of genomic DNA from 6-well plate and sanger sequencing. Clone 1D9 was further used for imaging.

In order to generate a control cell line allowing to measure experimental uncertainty on green-red distances, a clone previously engineered (1F11, Mach et al.) containing a TetO and LacO inserted on chromosome 15 separated by 150kb, but not expressing any fluorescent repressors, was used. 500'000 cells were transfected with PBase and two piggyBac vectors containing TetR-tdTomato and TetR-eGFP using Lipofectamine300 using 200ng of each plasmids. The media was changed 24h after transfection and cells were FACS sorted onto 96-well plates containing culture media supplemented with primocin antibiotics (InvivoGen, ant-pm-1). Clones were selected on the microscope for comparable signal to noise in both channels and compared to the dual operator lines.Clone 1B2 was then selected for imaging.

**Microscope setup**

Cells were imaged with a Nikon Eclipse Ti2 inverted widefield microscope equipped with a Total Internal Reflection Microscopy iLAS2 module (Gataca systems) for azimuthal illumination (*2*), a Perfect Focus System (Nikon) and motorized Z-Piezo stage (ASI) using a CFI APO TIRF 100×, 1.49NA oil immersion objective (Nikon). The microscope was operated in oblique illumination mode with a nominal incidence angle of 54 degrees. Excitation sources were a 473nm and a 552nm Omicron laserbench laser. Cells were maintained at 37 °C and 8% $CO_2$ using an enclosed microscope environmental control setup (Cube) and CO2 control (Brick) from Life science instruments. The microscope was equipped with two dual back-illuminated 95% quantum efficiency scientific CMOS cameras (Orca-fusion-BT, Hamamatsu) with a pixel size of 6.5 *µm*. A dichroic long-pass (550, Chroma) was added to separate red from green to the two cameras. Furthermore, two bandpass (514/44, Semrock, 595/65, Chroma, respectively) filters were added to the green and red channel.

**Live-cell image acquisition**

The microscope was controlled using NIS-Elements software (Nikon, version 5.42.07). Laser power was set to 1.5% of the maximum power for the 473nm and 552nm lasers which approximately corresponds to powers of 231 µW and 345 µW at the objective. The power was measured weekly using a power meter (Thorlabs, PM400) and a Silicon Photodiode (Thorlabs, S170C). The two cameras were aligned manually using TetraSpeck beads (100nm, Thermo Fisher Scientific, T7279) and finer correction was performed using the data collected as described below ('Correction of chromatic aberrations'). The field of view was binned 2 × 2 in the x and y directions which resulted in an effective pixel size of 130nm. Images were acquired for 15 minutes at 0.5 Hz (2 seconds between frames) and a Z-step size of 0.3 *µm* for 9 stacks. The total displacement of the Z-stage was 2.4 *µm* and the total time of acquisition was 15 minutes. Acquisition of the control cell line (1B2) was performed at 10% laser power in both channels to visually match signal-to-noise levels observed in the 1A2 cell line. The exact number of movies and tracks can be found in **Supplementary Table 2**.

**Fixed-cell image acquisition**

For fixed cell measurements to estimate the localization error, $10^6$ cells were seeded onto Mattek dishes and incubated for 24 h at 37 °C, 8% CO2. The medium was removed and the cells were fixed in 4% paraformaldehyde (Electron Microscopy Sciences, 15710) in PBS for 20 min at room temperature. The cells were washed three times in PBS and Fluorobrite medium was added to the Mattek dish to achieve comparable background fluorescence levels. The laser power in the red channel (552nm) was increased to 5% of the maximum power to achieve a comparable signal to noise ratio as in live cells. Images were acquired as described for live-cell image acquisition above.

### Image processing and construction of LacO-TetO distance tracks

Images were saved in the NIS-Elements format (.nd2) and processed in Python using the nd2 library available on GitHub. Due to the large size of the images, image analysis was parallelized using the Dask library. The imaging analysis pipeline was written in Python using the Snakemake library (*3*) for workflow management, and is available on GitHub at https://github.com/NessLfy/localization_precision_estimation.

#### *Nuclei segmentation*

Nuclei were segmented using the Stardist cell segmentation algorithm (*4*) with the pretrained "2D_versatile_fluo" model. Since the nuclei were not independently stained, segmentation was performed in the 552 nm channel on the maximum-intensity projection of the nucleoplasmic TetR-TdTomato signal. Segmentation was performed at each individual time point, and the resulting masks were converted to .zarr format using the *zarr_tools* (https://github.com/BaroudLab/zarr-tools) library for efficient storage and processing.

#### *Tracking of nuclei*

Nuclear segmentation yielded images with labeled nuclei. We tracked the nuclei using a custom algorithm that matches cells based on the distance between the centroids of segmented nuclei in consecutive frames. In short, a nucleus in frame *n* is assigned to the closest nucleus in frame *n+1* only if the relationship is reciprocal—that is, if nucleus A in frame *n* is closest to nucleus B in frame *n+1* and vice versa.

#### *Spot detection*

Raw 3D images in each channel were first preprocessed with a low-pass filter (preprocessing.lowpass from *trackpy* (https://doi.org/10.5281/zenodo.11522100), with σ = 1) to reduce high-frequency noise. The filtered images were then processed with a white top-hat filter (white_tophat from *skimage.morphology*), using a cylindrical structuring element with a radius of 2 pixels and a height of 9 z-slices. To minimize biases arising from uneven intensity distributions across the field of view, subsequent analysis was performed on a per-nucleus basis.

Spots were identified using a custom local maxima detection algorithm. A voxel was considered a local maximum if its intensity matched the maximum within a spherical neighborhood of radius 7 pixels. The detected local maxima were ranked by intensity, and the top five were retained as candidate spots.

Each candidate spot was then fitted to a 3D Gaussian function of the form:

$$A\, e^{-(x-x_0)^2/(2\,\sigma^2{}_{xy})+(y-y_0)^2/(2\,\sigma^2{}_{xy})+(z-z_0)^2/(2\,\sigma^2{}_{z})} + \text{offset}$$

where *A* is the amplitude, $\sigma_{xy}$ and $\sigma_z$ are the standard deviations in the lateral and axial directions, respectively, and $(x_0, y_0, z_0)$ denotes the Gaussian center, providing sub-pixel spot localization. Fitting was performed using the curve_fit function from scipy.optimize (*5*).

#### *Tracking of arrays*

Candidate tracks were generated by selecting the highest-intensity spot from each channel at every time point. Based on previously measured frame-to-frame displacements (at 30s frame rates) and spatial distances between the operators (*1*), tracks were retained only if the following conditions were met for more than 80% of the frames:

- The displacement between consecutive frames in each channel was less than 1 μm.
- The distance between corresponding spots in the two channels was less than 1.5 μm.

Tracks that did not meet these criteria were excluded from further analysis. For tracks that passed the filters, we refined the frames that initially failed by evaluating the remaining four candidate spots in both channels. We computed the distances between all pairs of candidate spots and selected the closest pair. If this new pair met the original criteria, it was added to the track; otherwise, the frame was marked as a gap.

***Exclusion of doublet signals***

We selected *bona fide* G1 (unreplicated) cells by assessing the presence of doublet signals from any of the two arrays, which is indicative of a replicated allele . By visual inspection of cells with replicated arrays, we noticed that they were systematically closer than 1*μ*m. Thus for each nucleus and frame, we measured the distances between the first and second highest intensity pixels in a spherical radius of XX pixels. We excluded a nucleus from analysis if the proportion of time points where the distance was less than 1 *μm* was larger than a threshold of 16%, which qualitatively corresponds to the proportion of tracks where the two most intense pixels are part of the same singlet signal. As further validation for the selected *bona fide* G1 nuclei, we manually inspected a subset of time frames from all movies and discarded any remaining nuclei presenting doublets.

***Correction of chromatic aberrations***

Because the two arrays were imaged in different channels (green and red), the measured distances between them were affected by chromatic aberration, due to wavelength-dependent refraction in the optical system. We corrected for these aberrations following the procedure described in Ref. (*6*). This approach exploits the fact that, in the absence of aberrations, the components of the distance vectors between the two spots should be symmetrically distributed around zero, as there is no spatial bias favoring one configuration over another. Chromatic aberrations introduce a systematic bias that shifts the mean away from zero and distorts the symmetry of the distribution (**Materials and Methods Fig. 1**, right panels).

To correct this bias, as in Ref. (*6*), for each day of acquisition we fit a linear regression model to each coordinate component (x, y, z) of the distance vectors and extract a 3×3 correction matrix. This matrix is then applied to compensate for the bias introduced by chromatic aberrations.

Additionally, on each acquisition day we imaged 3D stacks of TetraSpeck beads (100 nm, Thermo Fisher Scientific, T7279) to validate the correction. This control confirmed that the linear model effectively corrected chromatic aberrations (**Materials and Methods Fig. 1**, top two panels). Putting together all the acquisitions of beads across days, the standard deviations of the X and Y components of the distance vectors between the green and red signals of the beads were reduced to 9 nm and 10 nm, respectively. The mean distance between the two colors decreased from 112 nm before correction to 10 nm after correction.

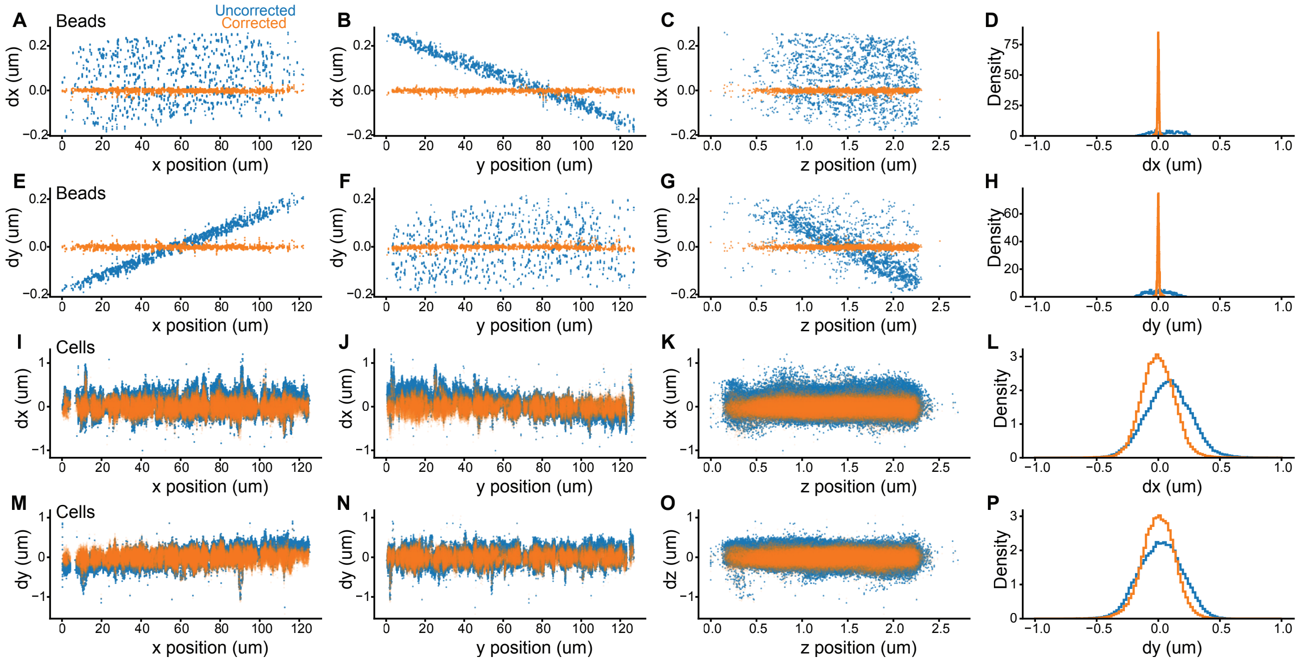


**Materials and Methods Figure 1**: Correction of chromatic aberration for one representative acquisition day. Top two panels: TetraSpeck beads; bottom panels: cells. **A-C.** X-component of the distance vectors of the beads between the red and green channels (dx) against the X, Y, and Z positions of the beads in the green channel before (blue) and after (orange) chromatic aberration correction. **D.** Distribution of dx before (blue) and after (orange) correction. **E-G.** Y-component of the distance vectors of the beads between the red and green channels (dy) against the X, Y, and Z positions of the beads in the green channel before (blue) and after (orange) chromatic aberration correction. **H.** Distribution of dy before (blue) and after (orange) correction. **H-K.** X-component of the distance vectors between the red and green operators (dx) against the X, Y, and Z positions of the green operators before (blue) and after (orange) chromatic aberration correction. **L.** Distribution of dx before (blue) and after (orange) correction. **M-O.** Y-component of the distance vectors between the red and green operators (dy) against the X, Y, and Z positions of the green operators before (blue) and after (orange) chromatic aberration correction. **P.** Distribution of dy before (blue) and after (orange) correction.

### Polymer model and simulations

***Polymer model***. The chromatin fiber is modeled as an unknotted polymer ring consisting of 4000 monomers, with each monomer representing 1 kilobase (kb) of chromatin. We define the unit of length as σ and the unit of temperature as T. By denoting $k_B$ the Boltzmann constant, the energy unit becomes ϵ = $k_B$T. All units are expressed in terms of these reduced units. Polymer connectivity is maintained via harmonic bonds, described by the potential $U_{bond}$, which connects consecutive monomers along the chain:

$$U_{\rm bond}(r) = K(r - r_0)^2$$

where $K = 100\ \epsilon/\sigma^2$ is the bond stiffness, and $r_0 = 0.8\ \sigma$ is the equilibrium bond length. To enforce bending rigidity, an angular potential $U_{angle}$ is applied between consecutive bonds:

$$U_{\text{angle}}(\theta) = K_{\text{angle}}(\theta - \theta_0)^2$$

where $K_{angle} = 0.5\epsilon$ is the angular stiffness, θ is the angle between two consecutive bonds, and $\theta_0 = 0$ is the equilibrium angle. This potential promotes slight alignment of neighboring bonds. Excluded volume interactions between monomers is simulated using a truncated and shifted Lennard-Jones (LJ) potential:

$$U_{\text{LJ}}(r) = \begin{cases} 4\epsilon \left[ \left(\frac{\sigma}{r}\right)^{12} - \left(\frac{\sigma}{r}\right)^{6} + \frac{1}{4} \right], & r \leq 2^{1/6}\sigma \\ 0, & r > 2^{1/6}\sigma \end{cases}$$

Finally, the polymer ring is simulated at a monomer volume concentration of $\rho\sigma^3 = 0.3$ in periodic boundary conditions.

***Simulation of loop extrusion.*** We simulate the one-dimensional loop extrusion process of cohesin using the Gillespie algorithm. The algorithm simulates the stochastic 1) binding, 2) unbinding, 3) stepping of cohesin. Given the rates associated with each of these processes, the Gillespie algorithm generates trajectories consistent with the underlying stochastic dynamics. Specifically, the algorithm assumes that the waiting time between successive events follows an exponential distribution, meaning that the temporal behavior of binding, unbinding, and stepping events is fully determined by their respective rates.

Cohesin is modeled as a bidirectional extruder, composed of two subunits that move in opposite directions along the polymer. In our simulations, we consider two scenarios for cohesin–cohesin encounters: (i) cohesins bypass one another, or (ii) they stall upon contact. Throughout the text, we explicitly state which of these conditions applies to each set of results. Cohesin stalling at CTCF sites is modeled in two ways: either as a complete barrier that prevents passage, or as a localized reduction in the stepping rate. The specific implementation used is always detailed alongside the corresponding results.

Finally, the coupling between the extrusion dynamics and the three-dimensional polymer simulation is achieved by representing extruders as additional harmonic bonds that connect the monomers occupied by the two cohesin subunits. These bonds are updated dynamically, based on the one-dimensional stochastic trajectories generated by the Gillespie algorithm.

***Simulation details.*** As specified before, we consider one chain consisting of 4000 monomers at a fixed concentration $\rho\sigma^3 = 0.3$ in a cubic box with periodic boundary conditions. The static and kinetic properties of the system are studied using fixed-volume and constant-temperature molecular dynamics (MD) simulations with implicit solvent and periodic boundary conditions. All MD simulations are performed using the LAMMPS package (7). The equations of motion are integrated using the velocity-Verlet algorithm. Time is expressed in MD units, where $\tau_{MD} = \sigma\sqrt{m/k_BT}$, and the integration time step is set to $\Delta t = 0.005\,\tau_{MD}$ to ensure numerical stability. The friction coefficient is set to $\gamma = 1/\tau_{MD}$.

For each system analyzed, we first equilibrate the polymer by integrating the equations of motion for $5 \times 10^8$to $1 \times 10^9$ time steps. After equilibration, we collect polymer configurations over an additional $5 \times 10^8$ time steps for production analysis. Each condition is simulated with 10 to 40 independent replicas.

### Supplementary Table 1: Oligonucleotides

| Oligonucleotides | Sequence (5' - 3') | Primer number |
|---|---|---|
| gRNA_KO_inside_TAD | CACGTTTGGCAACACCCTGC | |
| gRNA_100kb_inside_TAD | CTGCTACTACGTCCATCCAT | |
| geno_100kb_F | CGCACAAAGCCTCCTTG | 2354 |
| geno_KO_50-100kb_R | GACATGCTTGTGCTAGAGG | 2392 |

### Supplementary Table 2: Number of replicates

| Name of the cell line | number of movies | condition | number of tracks | number of unique data points |
|---|---|---|---|---|
| 1A2 (150 kb) | 213 | dTAG | 632 | 241574 |
| | 125 | WT | 786 | 300212 |
| | 74 | Fixed | 120 | 27175 |
| 1D9 (50 kb) | 287 | dTAG | 1269 | 479277 |
| | 192 | WT | 1196 | 447038 |
| 1B2 (control) | 460 | WT | 7484 | 7484 |
